# Global ribosome profiling reveals that mutant huntingtin stalls ribosomes and represses protein synthesis independent of fragile X mental retardation protein

**DOI:** 10.1101/629667

**Authors:** Mehdi Eshraghi, Pabalu Karunadharma, Juliana Blin, Neelam Shahani, Emiliano Ricci, Audrey Michel, Nicolai Urban, Nicole Galli, Sumitha Rajendra Rao, Manish Sharma, Katie Florescu, Srinivasa Subramaniam

## Abstract

The regulators that stall ribosome translocation are poorly understood. We find that polyglutamine-expanded mutant Huntingtin (mHtt), the Huntington’s disease (HD) causing protein, promotes ribosome stalling and physiologically suppresses protein synthesis. A comprehensive, genome-wide analysis of ribosome footprint profiling (Ribo-Seq) revealed widespread ribosome stalling on mRNA transcripts and a shift in the distribution of ribosomes toward the 5’ end, with single-codon unique pauses on selected mRNAs in HD cells. In Ribo-Seq, we found fragile X mental retardation protein (FMRP), a known regulator of ribosome stalling, translationally upregulated and it co-immunoprecipitated with mHtt in HD cells and postmortem brain. Depletion of FMRP gene, *Fmr1*, however, did not affect the mHtt-mediated suppression of protein synthesis or ribosome stalling in HD cells. Consistent with this, heterozygous deletion of *Fmr1* in Q175FDN-Het mouse model, Q175FDN-Het; *Fmr1^+/–^*, showed no discernable phenotype, but a subtle deficit in motor skill learning. On the other hand, depletion of mHtt, which binds directly to ribosomes in an RNase-sensitive manner, enhanced global protein synthesis, increased ribosome translocation and decreased stalling. This mechanistic knowledge advances our understanding of the inhibitory role of mHtt in ribosome translocation and may lead to novel target(s) identification and therapeutic approaches that modulate ribosome stalling in HD.

**One Sentence Summary:** Huntington’s disease (HD) protein, mHtt, binds to ribosomes and affects their translocation and promotes stalling independent of the fragile X mental retardation protein.

## INTRODUCTION

Huntingtin (Htt) is a ubiquitously expressed protein found throughout the nervous system and in non-neural tissues. Polyglutamine expansion (CAG expansion in Huntington disease gene homolog, *HTT)* results in mHTT, which causes Huntington’s disease (HD), a debilitative dominant brain disorder that affects people worldwide. Seminal studies have demonstrated the role of normal Htt in the development of mice. Mice deleted for Htt gene, *Hdh*, die around embryonic day 7.5–10.5, before the full emergence of the nervous system, indicating its role in cell survival and neurogenesis (1–4). Recent studies found that Htt deletion in the brain of younger mice (two months) led to death due to pancreatitis, and, in the adult brain, there was only a subtle defect in corticostriatal development and hyperactivity (5–7). The *HTT* gene, however, does not appear to affect development because HD human patients who are heterozygous or rare cases of homozygous for *HTT* appear to have normal development (8, 9). HD symptoms appear after birth, and the severity depends on the number of CAG repeats in *HTT*. Brain pathology and MRI studies show early damage to the striatum and cortex, and as the disease progresses, it affects multiple CNS and peripheral regions, leading to motor dysfunctions, weight loss, and energy deficits (20–22). These deficits in HD can emanate from one or more of the effects of mHtt on functions, such as vesicle- and microtubule-associated protein/organelle transport, transcription, autophagy as well as tissue maintenance, secretory pathways, and cell division (10–21). mRNA translation is crucial for cell growth and survival, and increasing data indicates that its dysregulations contributes to neurodegenerative disorders (22, 23). We previously identified mHtt as a potentiator of amino acid-induced mTORC1 signaling in association with Rheb and found that the selective upregulation of mTORC1 in the striatum worsened HD behavioral defects (24). However, the effect of mHtt on the regulation of protein synthesis remains less clear. Here, we investigated whether mHtt alters protein synthesis in both *in vitro* and *in vivo* models of HD and elucidated the mechanisms by which this might occur.

## RESULTS

### Suppression of protein synthesis and ribosome stalling in HD cells

We tested whether wtHtt/mHtt can alter protein synthesis by comparing the polysome profiles of striatal cells, which are immortalized neuronal cells isolated from the striatum of knock-in mice, containing a targeted insertion of a chimeric human–mouse exon 1 with 7 CAG (control), 7/111 CAG (HD-het) and 111/111 CAG repeats (HD-homo)(25). We observed a high polysome/monosome (PS/MS) ratio in the HD-homo cells compared with the HD-het or control cells (Fig. 1, A **and** B). We investigated whether the high PS/MS ratio in HD-homo cells is a reflection of more actively translating mRNA in HD-homo cells, using SUnSET, a non-radioactive, puromycin/antibody-based tool, to measure protein synthesis (26). We found diminished puromycin incorporation, reflective of reduced mRNA translation, in the HD-homo and HD-het cells compared with control cells (Fig. 1C). Puromycin incorporation was decreased by ∼20% and ∼40% in the HD-het and HD-homo cells, respectively (Fig. 1D). Like the puromycin, the incorporation of radiolabeled [^35^S]-methionine into newly synthesized proteins was also diminished by ∼40% in the HD-homo cells compared with control cells (fig. S1, A **and** B). A significant reduction in protein synthesis was also noted in human HD-het fibroblasts compared with unaffected controls (Fig. 1, E **and** F). Next, we asked why the PS/MS ratio was high in HD-homo striatal cells even though the protein synthesis was diminished. We hypothesized that the ribosomes paused, or elongate slowly, in the HD-homo compared with control striatal cells. To investigate this, we measured the translocation speeds of ribosomes in the HD-homo vs. the control striatal cells (ribosome run-off assay) with harringtonine, which blocks the initiation of mRNA translation without affecting ribosomes downstream (27, 28). Figure 1G shows a bigger PS in HD-homo striatal cells, compared to control striatal cells, at basal condition (Inset g1). A rapid increase in the 80S (monosome, MS) peak in control striatal cells, observed two minutes after treatment with harringtonine, compared with the HD-homo striatal cells (green arrow), indicating ribosomes run faster in control cells compared to HD. At this point, the PSs in the HD-homo cells remained high compared with those in control cells (Inset g2; red arrow) and, over time, there was a significant correlation after the harringtonine treatment (Fig. 1H), indicative of slowly moving ribosomes in HD. Within five minutes of the harringtonine treatment, the ribosomes appeared to complete the translocation both in the HD-homo and the control cells (Inset g3). We further confirmed that the HD-cells, despite the high PS/MS ratio, showed diminished protein synthesis under the harringtonine treatment (Fig. 1I). Collectively, these data indicated that the diminished protein synthesis in HD striatal cells is attributable to the slow elongation or stalling of the ribosomes compared with controls.

**Figure. 1.**
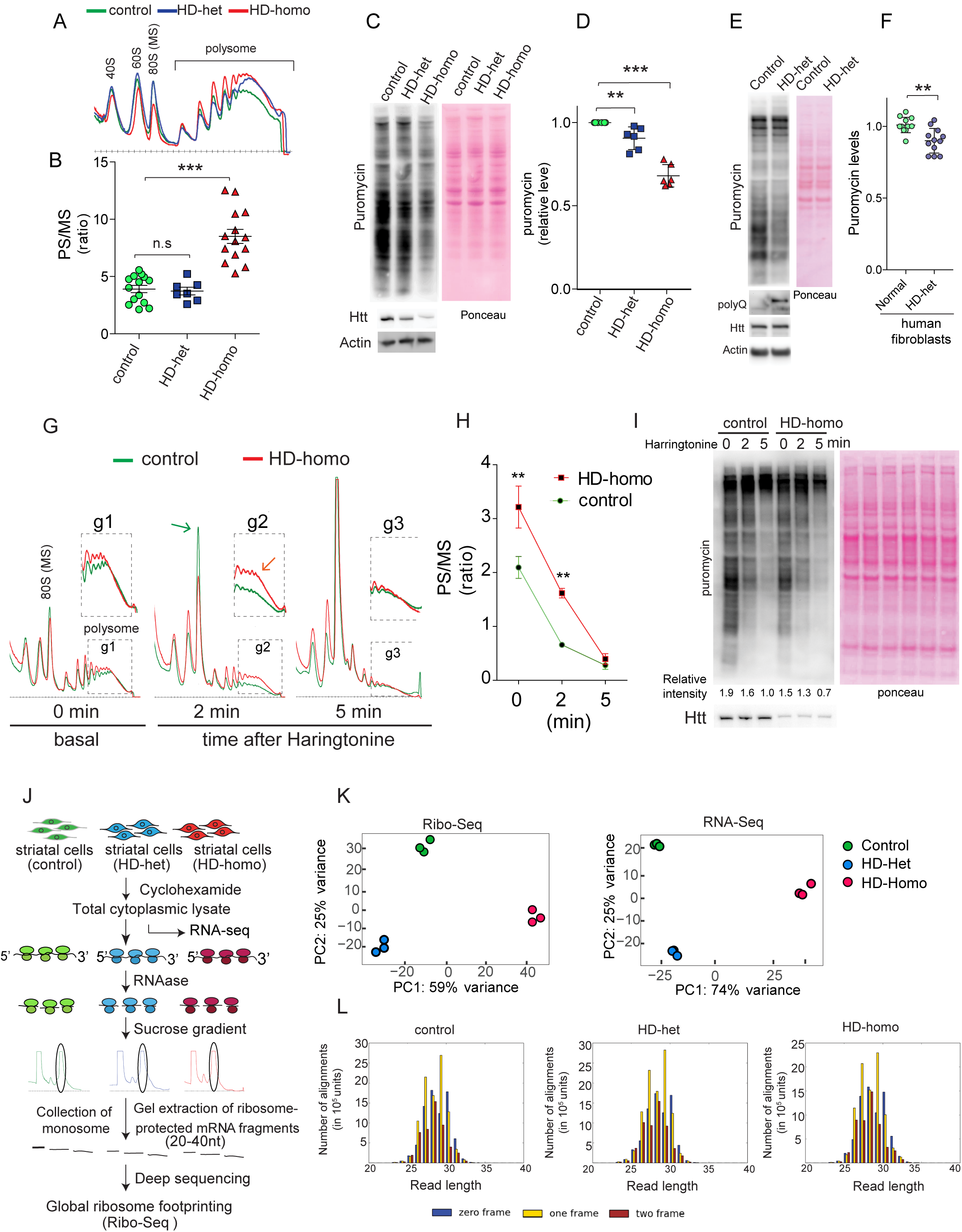
Suppression of protein synthesis and ribosome stalling in HD cells. (A) Representative polysome profiles of control and HD-het and HD-homo striatal cells. (B) Quantification of polysome to monosome (PS/MS) ratio in polysome profiles from A (using the undercurve areas). \*\*\**P* < 0.001 by Student’s *t* test. (C) Representative immunoblots of metabolic labeling of protein synthesis, using puromycin, and it quantification (D) in mouse striatal cells. Ponceau staining of the blots was used to quantify total protein signal in each lane. \*\*\**P* < 0.001 and \*\**P* < 0.01 by Student’s *t* test. (E) Puromycin metabolic labeling in human fibroblasts and (F) its quantification. \*\**P* < 0.01 by Student’s *t* test. (G) Representative polysome profiles obtained from mouse striatal cells in control and HD-homo stratal cells after ribosome run-off assay. Insets g1-g3 show ribosome movement between control (green) an HD-homo (red) cells. (H) Quantification of polysome to monosome (PS/MS) ratio in polysome profiles from G (using the undercurve areas). \*\**P* < 0.01 by Student’s *t* test. (I) Representative immunoblots of metabolic labeling of protein synthesis in control and HD-homo striatal cells after ribosome run-off assay. The numbers are representing the normalized densities of puromycin signals of each lane. (J) A schematic diagram showing the experimental design for performing the ribosome profiling (Ribo-Seq) and mRNA sequencing (RNA-Seq) in indicated mouse striatal cells. (K) Principal component analysis was performed using Ribo-Seq and RNA-Seq data obtained from control, HD-het and HD-homo striatal cells. (L) Representative graphs showing triplet periodicity plots generated for protein coding regions of Ribo-Seq data obtained from control, HD-het and HD-homo, striatal cells.

### Global ribosome profiling reveals diverse ribosome occupancy on mRNA transcripts in HD cells

To identify the mRNA transcripts on which ribosomes are stalled in HD cells, we employed ribosome profiling, a high-throughput sequencing tool that measures ribosome occupancy by sequencing ribosome-protected mRNA fragments (RPF) at a global translatome level (29–32). Figure 1J shows the research design, where three replicates of wild-type control striatal cells and HD-het and HD-homo mutant striatal cells were subjected to ribosome profiling in which we prepared ribosome footprints (Ribo-Seq) and matching RNA (RNA-Seq). These were then converted into sequenceable libraries and sequenced with deep coverage (see details in the Materials and Methods Section). Multiple quality control measures, such as principal component (Fig. 1K) and Euclidian distance analyses (fig. S2), showed that the replicates were very similar, with genotype as the principle source of differences between the control and the HD cells. Most of the RPF were mapped to annotated protein-coding ORFs (fig. S3A), with a 29-nt expected triplet nucleotide periodicity (Fig. 1L) and ribosome occupancy at the start and stop codon (fig. S3B). Thus, we have generated a high-quality Ribo-Seq library of control, HD-het and HD-homo, striatal cells.

**Figure 2.**
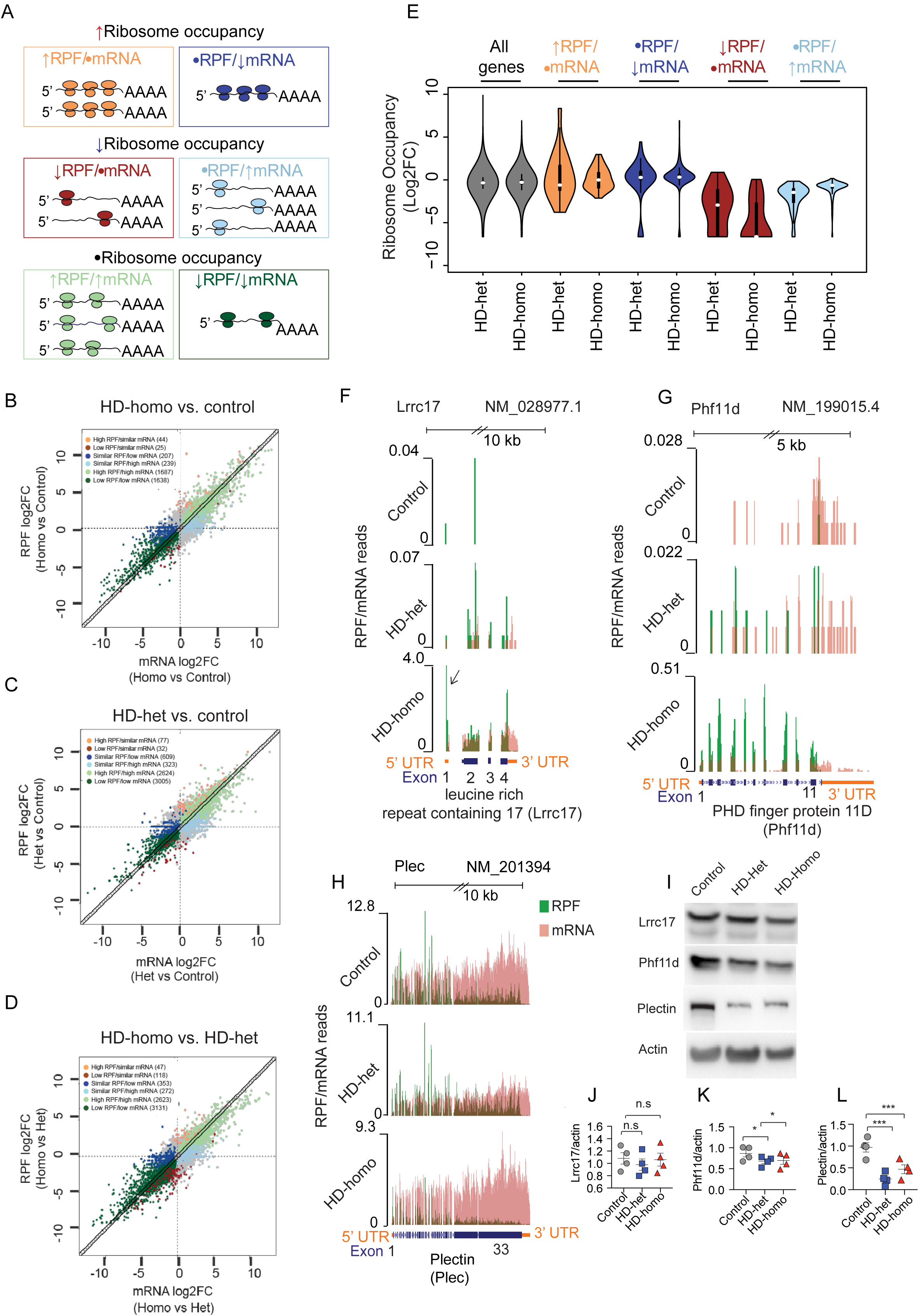
Global ribosome profiling reveals diverse ribosome occupancy on mRNA transcripts in HD cells. (A) A diagram showing how the changes in mRNA-Seq data were used to interpret Ribo-Seq data about ribosome occupancy. RPF; ribosome protected fragments. ↑; increased, ↓; decreased, and •; no change. (B, C, D) Scatter plots of expression changes of Ribo-Seq vs mRNA-Seq data in control, HD-het and HD-homo, striatal cells. As shown in A, changes in Ribo-Seq are classified into six groups. The numbers of genes in each group are shown at the top left corner of each plot. Total number of genes:36411, absolute fold change (FC) >2, nominal *P* <0.05, and false discovery rate (FDR) =0.15. (E) Plots showing the distribution of ribosome occupancy changes (calculated by number of Ribo-Seq reads divided by mRNA-Seq reads for each gene) in HD-homo and HD-het cells (compared to control cells). (F-H) Graphs showing overlay of Ribo-Seq/mRNA-Seq reads for indicated genes obtained from UCSC browser. (I) Representative immunoblots showing the protein expression levels of indicated gens in *F-H* and (J-L) the corresponding quantifications. \**P* < 0.05 and \*\*\**P* < 0.001 by Student’s *t* test.

As has been described before (33), several relationships between RPF and mRNA can be deciphered using ribosome profiling data. Because ribosomes are stalled in HD cells (Fig. 1A-I), our goal was to identify mRNA targets with high RPF in HD. For this, we used Anota2Seq software (34), which applies partial and random variance models to analyze differential RPF (a.k.a. translation efficiency) between control and HD cells to estimate the changes within each RNA source (RPF or mRNA) (35). Figure 2A shows the six categories of RPF/mRNA groups, showing differential ribosome occupancy, between control, HD-het, and HD-homo, striatal cells, that were expected in our analysis, listed as follows: a) high RPF/similar mRNA (↑RPF/•mRNA), b) similar RPF/low RNA (•RPF/↓mRNA), c) low RPF/similar mRNA (↓RPF/•mRNA), d) similar RPF/high mRNA (•RPF/↑mRNA), e) high RPF/high mRNA (↑RPF/↑mRNA), and f) low RPF/low mRNA (↓RPF/↓mRNA). As shown in the scatterplot (Fig. 2, B-D) among 36,411 genes, 3,840 mRNA targets in the HD-homo cells, and 6,670 mRNA targets in the HD-het cells showed significantly changed (p-value <0.05) ribosome occupancy (RPF/mRNA) compared with controls (Fig. 2, B **and** C; **Data files S1, S2**), and 24.6% (2,074) with high RPF were similar between the HD-homo and HD-het cells (**Data file S3**). Differential RPF/mRNA was also observed between the HD-homo and HD-het cells (Fig. 2D; **Data file S4)**. In the RPF categories high RPF/similar RNA and similar RPF/low RNA, which reflect high ribosome occupancy normalized to mRNA, we found 251 transcripts in the HD-homo and 686 transcripts in the HD-het cells compared with controls. Robust differences found in the RPF of the other categories of HD (Fig. 2, B **and** D) might indicate transcript level changes consistent with the transcriptional role of mHtt (36). Furthermore, we confirmed that the Anota2Seq algorithms were able to separate the ribosome occupancy of each category from HD-het and HD-homo cells compared with controls (Fig. 2E). Gene ontology (GO), using the Ingenuity Pathway Analysis (IPA) for each category, showed that the genes belong to diverse signaling pathways, including IL-10 signaling, Alzheimer’s disease signaling, AMPK signaling, cAMP signaling, Rho GTPase signaling, matrix metalloprotease pathways and EIF2, and mTOR (fig. S4). We examined the ribosome profiles of some of the targets in the Ribo-Seq and their protein expressions. We combined the uploaded profiles, as a track hub in the University of California Santa Cruz (UCSC) Genome Browser (37), from the triplicate experiments and overlaid selected RPF (green) normalized to corresponding mRNA (orange) (Fig. 2, F-H). The leucine-rich repeat-containing protein 17 (Lrrc17) showed few reads in the control but higher ribosome density across the protein-coding region in HD-homo compared with HD-het; there was also an even higher ribosome density in the HD-homo 5’ UTR region (arrow) (Fig. 2F). However, at the protein level, we did not see any difference in Lrrc17 between the groups, suggesting that Lrrc17 might be regulated via protein stability (Fig. 2, I **and** J). Phf11d, a PHD zinc finger protein consisting of 11 exons, showed a varying degree of high RPF on exons 2–10 in the HD-homo and HD-het cells compared with controls (Fig. 2G), and unlike Lrrc17, its protein levels in the HD-homo and HD-het cells were significantly diminished compared with controls (Fig. 2, I **and** K). Plectin, an intermediate filament-binding protein, showed a varying degree of high RPF across its 33 exons, with an overall increase in RPF at exon 33 in the HD-het cells compared with controls (Fig. 2H), but its protein levels were dramatically diminished in HD-het and HD-homo striatal cells compared with a control (Fig. 2, I **and** L). Therefore, despite high RPF and mRNA density, these targets showed no increase in protein levels, which suggested that the ribosomes were potentially stalled on these mRNA transcripts and supported the ribosome run-off experiment (Fig. 1, G-I). Additional examples of overlaid selected RPF normalized to corresponding mRNA transcripts are shown in sfig 5 and described in the associated **supplementary text**. A full UCSC Genome Browser link for the global ribosome footprints pooling all three replicates for control, HD-het and HD-homo cells will be available soon.

**Figure 3.**
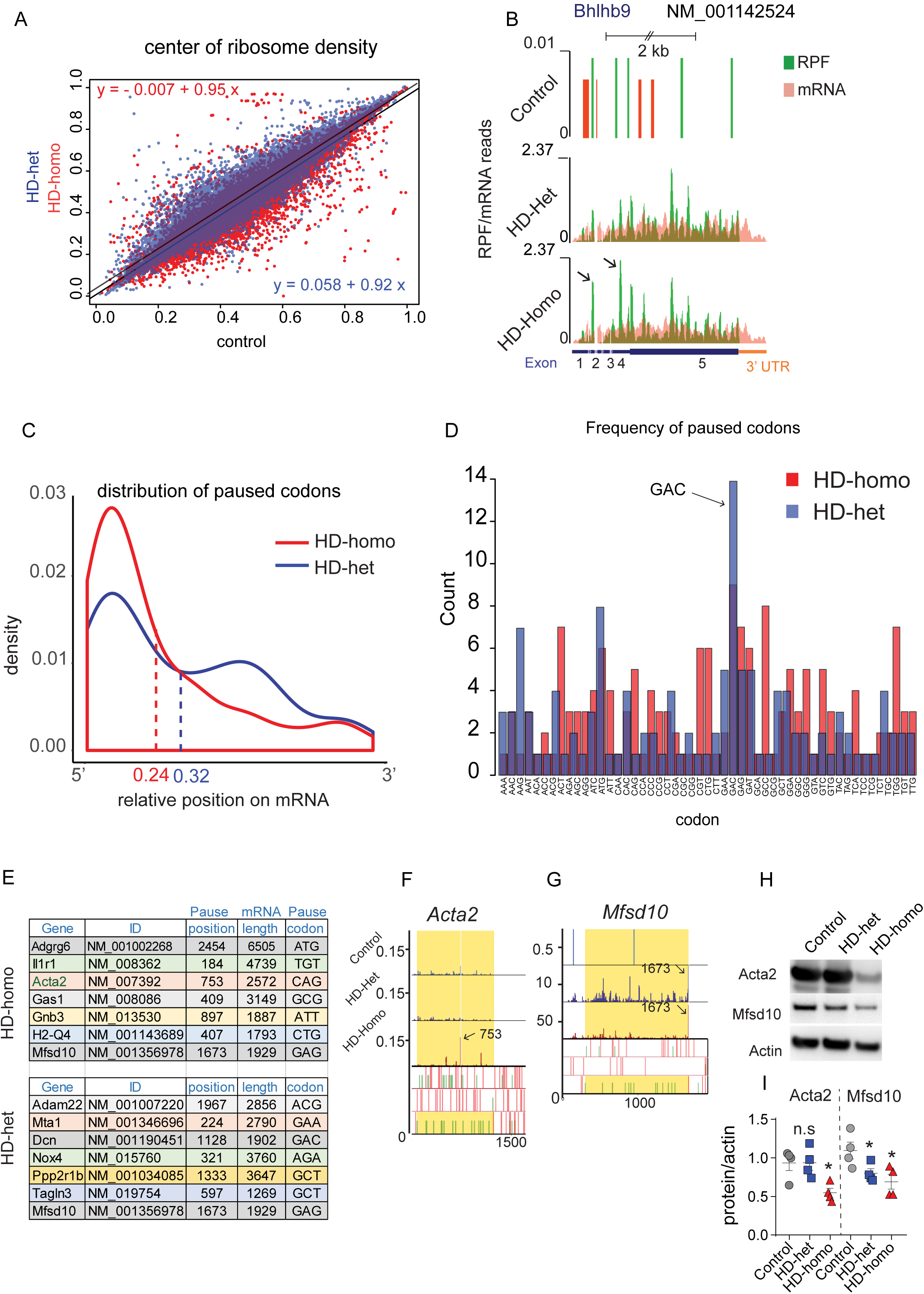
Widespread 5’ end ribosome occupancy and single codon pauses in HD cells. (A) Scatter plots showing the distribution of “center-weight analysis” score for each gene in HD-homo (red) and HD-het (blue) versus control cells. The diagonal lines represent the linear regression models and the calculated formulas are mentioned at the top left corner (HD-homo) and bottom right corner (HD-het) of the plot. (B) Graphs showing overlay of Ribo-Seq/mRNA-Seq reads for Bhlhb9 gene obtained from UCSC browser. (C) Pause density plot showing the distribution of common paused codons (found among 3 replicates of HD cells versus control cells) over their corresponding mRNAs. Numbers showed at the bottom represent the average of relative positions of paused codons in HD-homo (red) and HD-het (blue) cells. (D) Bar plots showing the frequencies of paused codons from C. (HD-homo cells; red, and HD-het cells; blue). (E) Tables representing top paused transcripts in HD-homo and HD-het cells. (F & G) Graphs representing codon pauses of the selected transcripts from E. Arrows are indicating the position of paused codons. (H) Representative immunoblots showing the protein expression levels of paused transcripts (from F & G) in HD-het and HD-homo cells and (I) corresponding quantifications. *P < 0.05 by Student’s t test.

**Figure 4.**
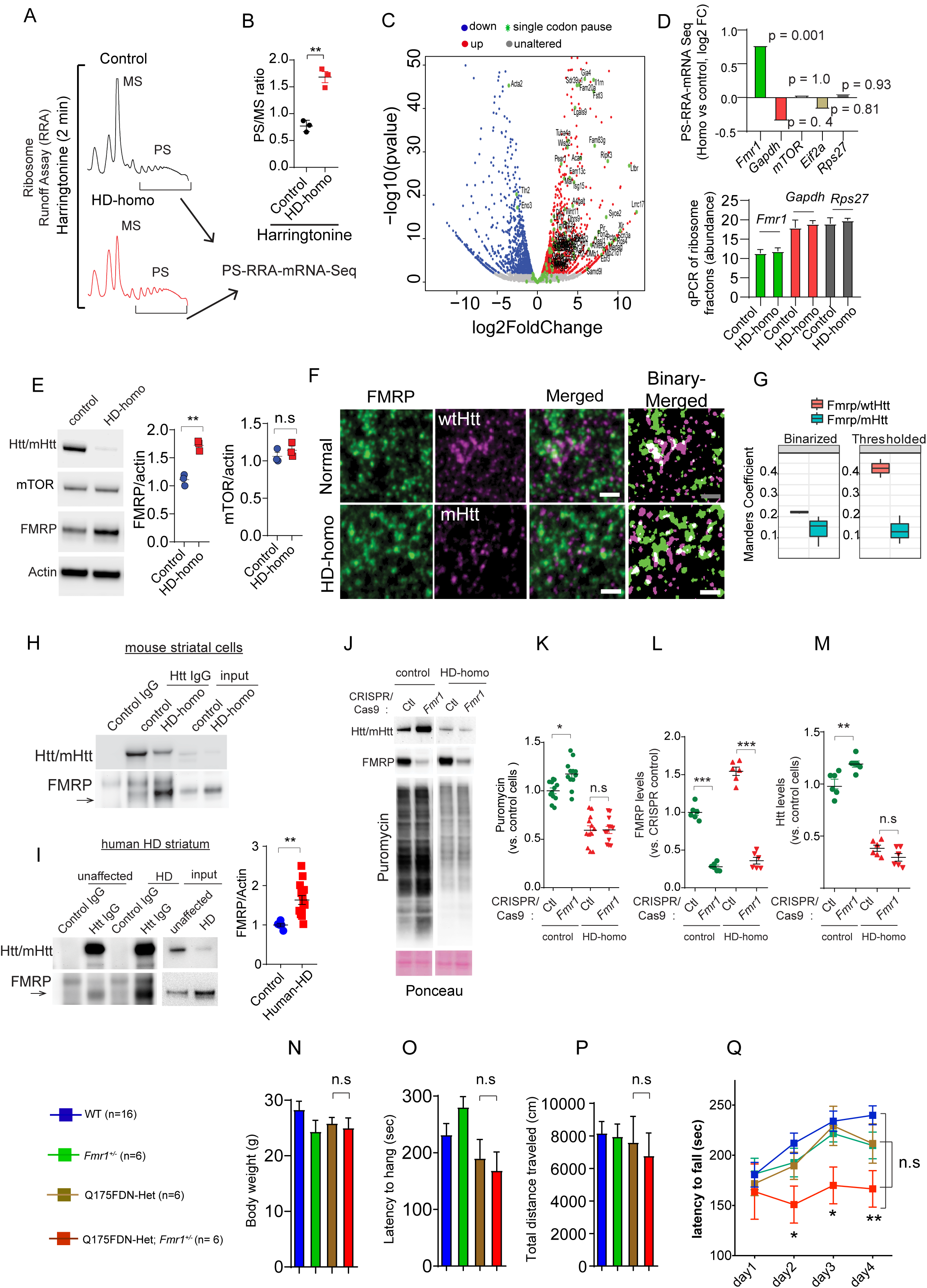
*Fmr1* depletion has no discernible effect on protein synthesis in HD cells, and in HD mice show subtle defects in motor learning skills. (A) Graphs representing polysome profiles of control and HD-homo cells after ribosome runoff assay. Arrow indicate mRNA-Seq was performed using RNAs isolated from polysome containing fractions. (B) Quantification of polysome to monosome (PS/MS) ratio in polysome profiles from A (using the undercurve areas). (C) Volcano plot representing changes in the expression levels of ribosome bound mRNAs in HD-homo cells (vs control cells) after ribosome run-off. mRNA transcripts with absolute log2 fold change > 0.3 and p value < 0.05 were shown with red (up, increased polysome binding), blue (down, decreased polysome binding), grey (unaltered, no difference in polysome binding), or green asterix, representing detected paused genes by Ribo-Seq in HD-homo cells (from Figure 3-C). (D) A bar plot representing mRNA levels of some known translator regulator genes within polysome fractions obtained from HD-homo cells (comparing to controls) after ribosome runoff assay (upper panel), and qPCR of the indicated targets in the purified ribosome fractions (bottom panel). (E) Representative blots and corresponding quantifications showing protein levels of Htt, mTOR and FMRP in control and HD-homo cells. **P < 0.01 by Student’s t test. (F) The colocalization between Htt (red) and FMRP (green) was measured in fixed cells with STED. (G) The Manders’ colocalization coefficients were calculated for the ratio of [FarRed-label] colocalizing with [Red-label], for both control and homo conditions, using background subtracted (thresholded) and binarized STED images. (H and I) Representative immunoblots of immunoprecipitation assays (using anti Htt antibody) on mouse striatal cells (H) and postmortem human striatal tissues (I, *left panel*). I, *right panel*, shows FMRP levels in unaffected and HD human post-mortem striatum. (J) Representative immunoblots performed on control and *FMR1* depleted mouse striatal cells after puromycin metabolic labeling. (K, L & M) Quantification of blots from J showing puromycin incorporation (K), FMRP levels (L), and Htt levels (M) in *Fmr1* depleted and control striatal cells. Bar plots representing the body weights (N), inverted hanging (O), open field (P), and rotarod (Q) experiments performed on ∼10 months old Q175FDN-Het; *Fmr1*+/– mice (comparing to wt, *Fmr1*+/- and Q175FDN-Het mice/equal sex ratio). \*\*\**P* < 0.001, \*\**P* < 0.01, by two-way ANOVA followed by post-hoc Bonferroni multiple comparison test.

**Figure 5.**
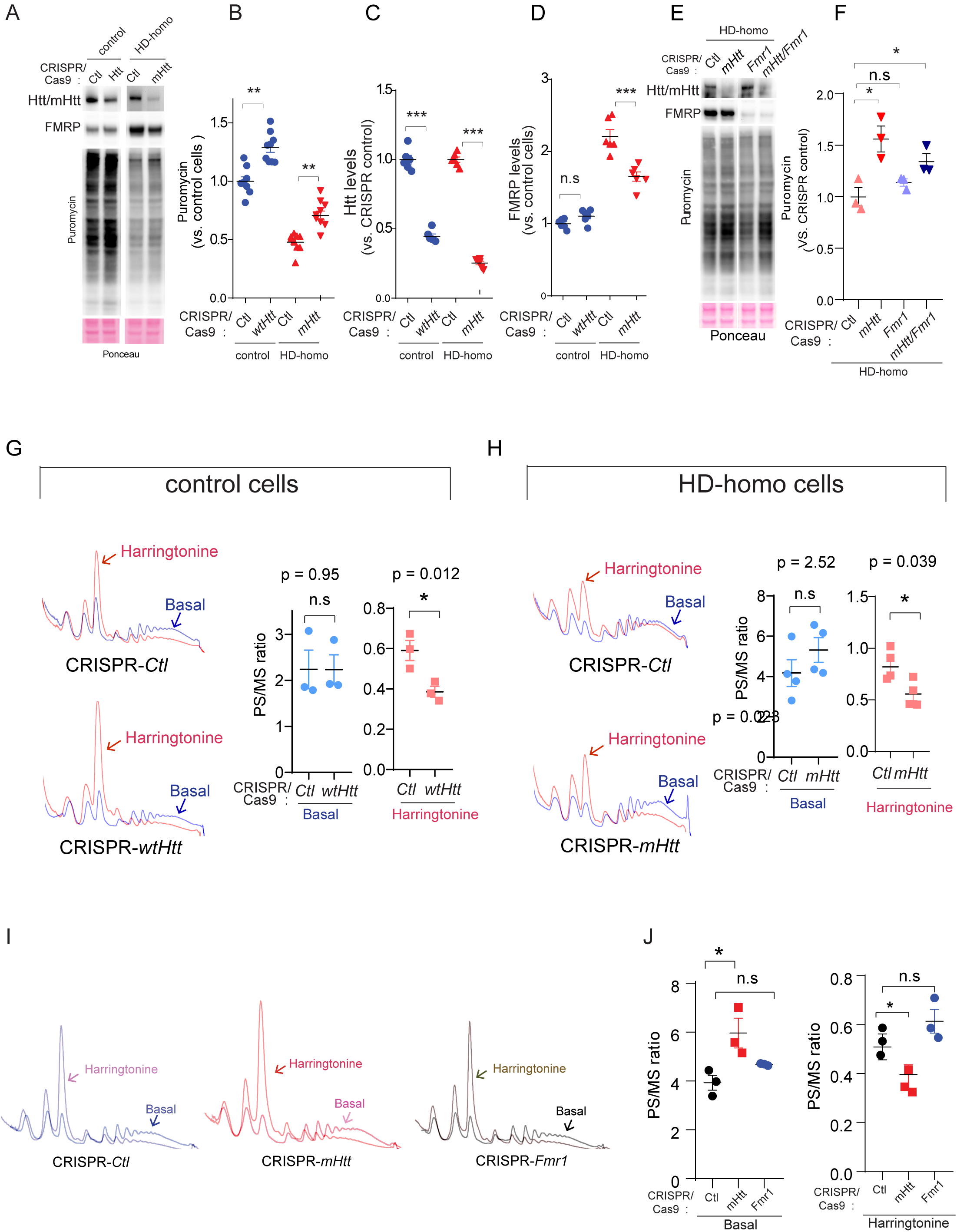
Depletion of wtHtt/mHtt enhances protein synthesis and increases the speed of ribosome translocation. (A) Representative immunoblots performed on Htt depleted mouse striatal cells after puromycin metabolic labeling. (B, C & D) quantifications of the blots from A showing puromycin incorporation (B), levels of Htt (C) and FMRP (D) in *Htt* depleted control, HD-homo striatal cells. (E) Representative immunoblots performed on *Htt* or *Fmr1* depleted, or *Htt/Fmr1* double depleted HD-homo cells after puromycin metabolic labeling. (F) quantifications of the blots from *P* showing puromycin incorporation. (H-Q, \*\*\**P* < 0.001, \*\**P* < 0.01, *P < 0.05 by Student’s *t* test). (G&H) Representative graphs showing polysome profiles obtained from *Htt* depleted mouse striatal cells before (basal) and after ribosome runoff assay using harringtonine (2 min) and their corresponding quantification of polysome to monosome (PS/MS) ratios (using undercurve areas). (I&J) Representative graphs showing polysome profiles obtained from *mHtt* or *Fmr1* depleted mouse striatal cells before (basal) and after ribosome runoff assay using harringtonine (3 min) and their corresponding quantification of polysome to monosome (PS/MS) ratio (using undercurve areas) *P < 0.05 by Student’s *t* test.

Altogether, for the first time, the super-resolution ribosome profiling in HD cells revealed a) enhanced ribosome occupancy but diminished protein expression for the selected targets, b) abundant transcript level RPF changes, and c) alterations in the cellular and molecular pathways, including the autism, apoptosis, and mTOR signaling pathways.

### 5’ end ribosome occupancy and single codon pauses in HD cells

Next, we employed PausePred software (38) to perform a ribosome pause analysis. For genes with detected pauses, by determining the center of the ribosome density (39) we were able to estimate whether there was any shift in the ribosome occupancy toward the 5’ end compared with the HD and control RPF data. We found 19, 498 mRNA transcripts in HD-homo and 16, 773 mRNA transcripts in HD-het in which the center of the ribosome density was shifted toward the 5’ end (Fig. 3A), compared with controls (**Data files S5 and S6**). We also observed that while some genes practically had no footprints in the controls, they had footprints in their Het/Homo profiles, which implies transcriptional changes (**Data files S7 & S8**). For example, Bhlhb9, basic helix-loop-helix transcription factor, contained high RPF/RNA in the HD-homo or HD-het cells but very few or no RPF/RNA in controls; even then, the RPF were concentrated toward the 5’ ends on exons 2 and 4 in the HD-homo cells compared with the HD-het cells (Fig. 3B; **arrow**).

Because the translocation of ribosomes occurs along the mRNA one codon at a time, we investigated the potential of single-codon resolution pauses on mRNA in HD cells compared with controls. Using PausePred software, which infers the locations of ribosome pauses (38), we analyzed global single codon pauses between the control and HD cells. We found numerous pauses between control and HD cells, but there were differences within the control and HD replicates (fig. S6), indicating that, in cells, the ribosome pauses on a given codon is not static but rather a dynamic event. We then pooled all triplicates and found ∼165 targets in the HD-homo cells and ∼125 targets in the HD-het cells that showed one or more codon-specific pauses, compared to control (**Data files S9** and **S10**). Most of the targets showed ribosome pauses toward the 5’ end of the coding mRNA, both in the HD-homo and HD-het cells (Fig 3C). The frequency of paused codons revealed no specific preferences, although approximately 15% of the hits showed a pause at GAC in the HD-het cells (Fig 3D; **arrow)**. Figure 3E indicates the pause positions, total length, and the codon of the selected mRNA in the HD-homo and HD-het cells. For example, Acta2, a member of the actin gene family, showed a pause at position 753 (CAG) in the HD-homo, and Mfsd10, an ion transporter, showed a single codon pause at position 1673 (GAG) in the HD-homo and HD-Het cells (Fig 3, E-G; **arrow**). At the protein level, we found that both Acta2 in the HD-homo and Mfsd10 in the HD-homo and HD-het cells were highly downregulated, compared to control (Fig. 3, H **and** I). In the HD-het cells, for example, Adam22, a catalytically inactive member of transmembrane ADAM metalloproteases linked to synaptic functions (40), was paused at position 1967 (ACG) (fig. S7A). Similarly, Adgrg6, an adhesion G protein-coupled receptor G6, which couples to G(i)-proteins and G(s)-proteins, showed a pause at 2454 (ATG) in the HD-homo cells (fig. S7B). Therefore, the PausePred analysis demonstrated a widespread global 5’ ribosome occupancy and single-codon pauses in selected mRNA targets in HD cells.

**Figure 6.**
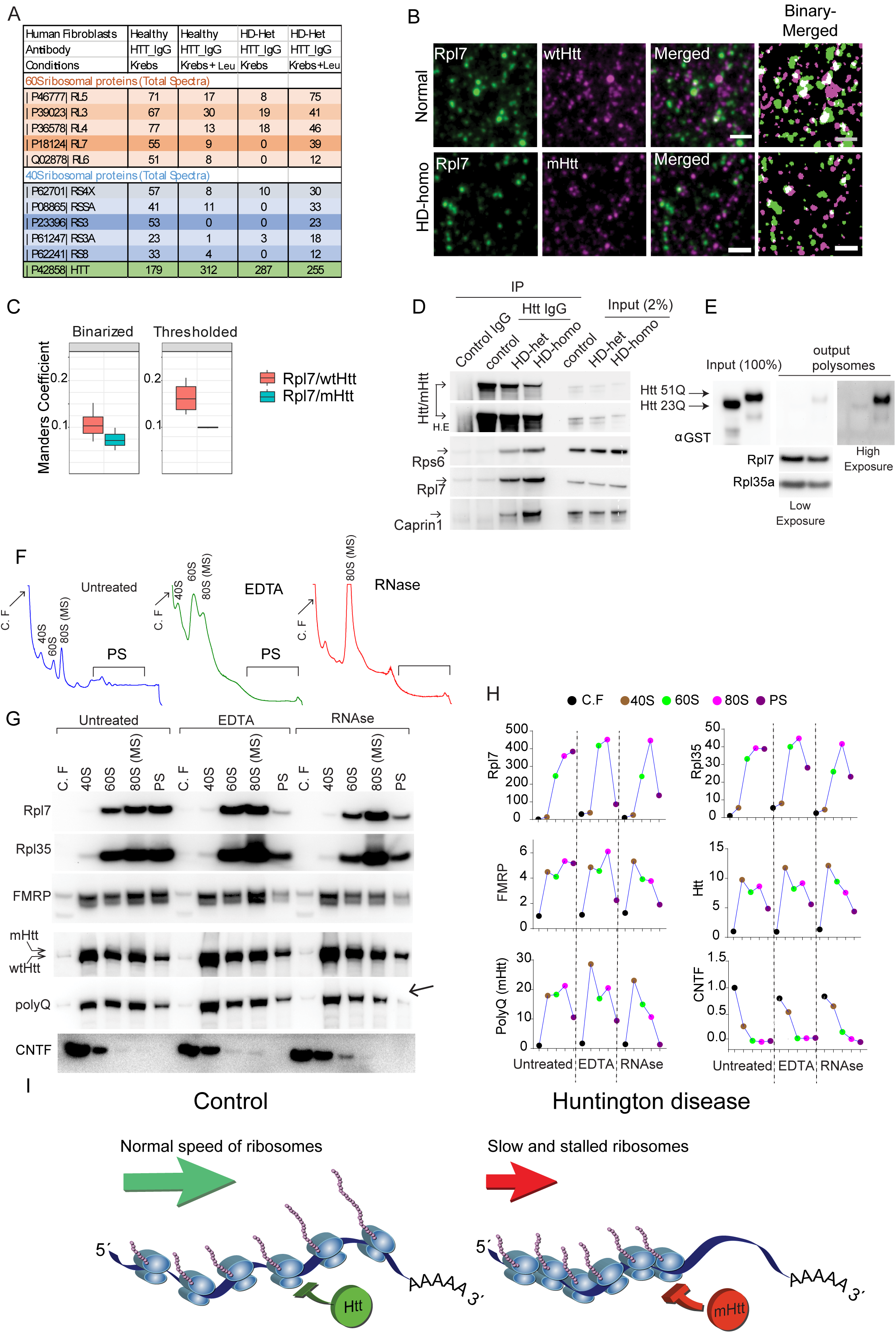
mHtt interacts directly with ribosomes in vitro and to an RNase-sensitive component of ribosomes in the brain. (A) Proteomics analysis on samples prepared from immunoprecipitation experiments on human (healthy controls 23Q^+/-^ and HD patient 69Q^+/-^) fibroblasts using an antibody against Htt. (B) Representative images of STED microscopy on mouse striatal cells using antibodies against Rpl7 (green) and Htt (red) counterstained with nuclear marker DAPI (see full image in SFig10B) (C) The Manders’ colocalization coefficients were calculated for the ratio of [FarRed-label] colocalizing with [Red-label], for both control and homo conditions, using background subtracted (thresholded) and binarized STED images. (D) Representative immunoblots showing co-immunoprecipitation of Htt/mHtt (MAB2166) with indicated proteins, and input. (E) Representative immunoblots on inputs and outputs of invitro ribosome binding assays using recombinant GST-exon1Htt 23Q and GST-exon1Htt 51Q proteins and isolated ribosomes from mouse striatal cells (see sfig 14B for the experimental diagram) and corresponding polysome profiles). (F) Representative graph showing polysome profile obtained from Q175FDN-Het mouse cortex under indicated conditions. (G) Representative immunoblotting experiments on polysome fractions obtained from Q175FDN-Het mice cortex. Note that all fractions containing polysomes were concentrated using a centrifugal filter unit and run as a single sample in immunoblotting. Also, an antibody against poly-Q was used to identify mHtt at the same location of Htt signal. Cntf serves as a negative control for ribosome binding protein. Cntf; ciliary neurotrophic factor, Rpl7; ribosomal protein L7, Rps6; ribosomal protein S6, Htt; huntingtin. (H) Schematic diagrams representing differences in proteins levels among different mouse cortex polysome fraction (from *G*). (I) Model showing wtHtt bind to ribosomes and physiologically inhibits the translocation of ribosomes, a normal function that is further enhanced by mHtt leading to slower movement and stalling of ribosomes.

### The identification of slowly translating mRNA reveals novel targets of ribosome pauses in HD cells

Diminished protein synthesis or slowly translating mRNA targets in HD cells can be among any category of RPF/mRNA (Fig. 2A-D). To identify the slowly translating mRNA in HD we took advantage of ribosome run-off assay. Because the ribosome run-off rate is slow in HD cells (Fig. 1, G **and** H), we hypothesized that identifying slowly translating mRNAs would further refine the targets of ribosome stalling in HD cells. To address this, we isolated mRNAs from the stalled polysomes (PSs) by employing a harringtonine-based ribosome run-off assay (RRA), followed by RNA-Seq (PS-RRA-mRNA-Seq) (Fig. 4A). Figure 4B shows the high PS/MS ratio obtained after the harringtonine treatment, consistent with Fig 1, G and H, indicating that ribosomes run slowly in HD cells compared with controls. There were ∼1157 targets that showed high mRNA abundance (red dots) and ∼1248 targets that showed low mRNA abundance (blue dots) in the PSs of HD-homo cells (p< 0.05) compared with control PSs in the PS-RRA-mRNA-Seq data (Fig. 4C; **Data file S11**). Next, we compared the single-codon paused targets (**Data file S9**) with the PS-RRA-mRNA-Seq. We found almost all the ∼130 single-codon paused targets in the highly abundant PS-RRA-mRNA-Seq hits (Fig. 4C; **indicated in green**). For example, the top PS-RRA-mRNA-Seq hit, Mfsd10 (paused at 1673) that had high RPF at the 5’ and 3’ ends (fig. S8A), and Acta2 (paused at 753) that had ∼15 fold less RPF in HD-homo compared to control (fig. S8B), showed diminished protein levels in the HD cells (Fig. 3, F-I). Therefore, the isolation of high/low abundance mRNAs from slowly elongating ribosomes revealed novel hits, and at least some of them we tested showed diminished protein levels, in HD cells.

### *Fmr1* depletion fails to rescue protein synthesis inhibition in HD cells

To decipher the mechanisms of ribosome stalling in HD cells, we looked for some of the known regulators of stalling in the PS-RRA-mRNA Seq data (**Data file S11**). The fragile X mental retardation protein (FMRP) encoded by *Fmr1*, a known promoter of ribosome stalling (41), was high in the RRA-RNA-Seq in the HD-homo cells (p = 0.001), but *Gapdh, mTOR, Eif2,* and *Rps27* were not significantly altered (Fig. 4D; **upper panel**). qPCR analysis of the ribosome fractions (MS and PS) not treated with harringtonine showed similar levels of *Fmr1*, *Gapdh*, and *Rps27* mRNA in both the control and HD cells (Fig. 4D; **bottom panel**). This indicates that *Fmr1* mRNA, which was found in high levels in the PS-RRA-mRNA-Seq, represents a likely target for slower moving ribosomes in HD cells. However, FMRP protein levels, but not mTOR levels, were upregulated in the HD-homo cells (Fig. 4E). Analysis of the RPF/RNA of *Fmr1* showed higher RPF accumulation on exons 5, 14, and 16 in the HD-homo cells, compared with the HD-het and controls (fig. S9A); however, both the HD-homo and HD-het cells had higher FMRP protein levels (fig. S9B), indicating that FMRP upregulation in HD cells might be a translational control by mHtt. One possibility would be that mHtt may directly interact with *Fmr1* mRNA and increase its translation. Consistent with notion, we found *Fmr1* mRNA bound predominantly with mHtt, but not wtHtt in an RNA immunoprecipitation-qPCR assay (data not shown). Another possibility would be that mHtt interact with FMRP bind and regulate mRNA translation in HD cells. Using super-resolution STED (stimulated emission depletion) microscopy we found that the endogenous wtHtt and mHtt colocalized with FMRP in immunohistochemically stained and fixated cells (Fig. 4F, **full image** fig. S10A). Using object-based colocalization methods, we found that around 25—35% of Htt clusters were in proximity (<300 nm) to the aforementioned proteins, with clear differences observable between the control and HD-homo conditions when calculating the Manders’ Colocalization Coefficients (Fig. 4G). Next, we tested FMRP and wtHtt/mHtt interaction by co-immunoprecipitation. We found that endogenous FMRP interacted strongly with mHtt in HD striatal cells (Fig. 4H), as well as in human HD striatum (Fig. 4I**, *left panel***), which, like HD-homo cells (Fig. 4E), also showed high FMRP protein levels compared to healthy controls (Fig. 4I**, *right panel*)**. Thus, FMRP protein is upregulated in HD and both protein and mRNA of FMRP interacts strongly with mHtt compared to wtHtt.

These data prompted us to hypothesize that FMRP might regulate ribosome stalling in HD cells. However, CRISPR/Cas-9-induced FMRP depletion significantly increased protein synthesis only in control cells but not in the HD-homo cells (Fig. 4, J **and** K). The depletion of FMRP was comparable (∼80%) between the control and the HD-homo cells (Fig. 4L). Intriguingly, the depletion of FMRP significantly elevated the levels of wtHtt but not mHtt (Fig. 4M), a finding consistent with a previous report that showed that Htt mRNA is a potential translationally downregulated target of FMRP (42). Together, these data suggest that although FMRP is upregulated in HD cells, it has no apparent effect on the inhibition of protein synthesis in HD cells. Therefore, we conclude that mHtt-mediated suppression of protein synthesis in HD cells is either upstream or independent of FMRP. Enhanced phosphorylation of Eif2α (pEif2α, S51), an indication of general stress response, can diminish protein synthesis (44), but we found reduced levels of Eif2α protein with no significant pEif2α alterations in HD-homo cells compared to control cells, (fig. S11A), consistent with a previous report (46). The ribosome occupancy of Eif2α (contains 14 exons) showed high RPF in exon 3 and 10, and where as Eif2α kinases, Perk (contains 17 exons) and Gcn2 (contains 39 exons), showed high ribosome occupancy towards 5’ region distributed on exons 1, and there was no discernible difference between control and HD-homo cells (fig. S11B-D). These data suggest that the inhibition of protein synthesis in HD is independent of FMRP and Eif2α signaling.

### *Fmr1* depletion in HD mice show no discernable phenotype, but promotes subtle deficits on motor learning skills

Although the cell culture data indicated that FMRP depletion had no significant effect on reversing protein synthesis inhibition, we wondered whether FMRP might regulate HD-linked behaviors *in vivo*. This notion was further strengthened by the data that showed that mHtt interacts with FMRP and is upregulated in HD cells (Fig. 4, E-I). Therefore, to address *in vivo* relevance of FMRP in HD, we generated HD mice, Q175FDN-Het (45), lacking one copy of the *Fmr1* gene (Q175FDN-Het; *Fmr1^+/–^*). At ∼10 months of age, we found no differences in body weight (Fig. 4N), muscle strength (Fig. 4O), or locomotion (Fig. 4P) in these mice compared with control groups (n = 6-16/equal sex ratio). But, while, the HD mice had no observable phenotype, consistent with a previous report (45), and acquired motor learning skills (rotarod) comparable to WT or *Fmr1^+/–^* mice, the Q175FDN-Het; *Fmr1^+/–^* mice showed a subtle defects to learn motor skills, compared to WT mice (Fig. 4Q). Thus, consistent with lack of FMRP depletion on protein synthesis in HD cells, the FMRP heterozygous mice have no gross effects on HD phenotype, but show a subtle deficit on motor learning, suggesting mHtt and FMRP may regulate the translation of a selected mRNAs involved in motor learning in the brain.

### Depletion of wtHtt or mHtt enhances protein synthesis and increases the speed of ribosome translocation

Next, we hypothesized that if mHtt blocks protein synthesis upstream, or independent, of FMRP, then the depletion of mHtt should increase protein synthesis. We found that the CRISPR/Cas-9 depletion of wtHtt or mHtt robustly increased protein synthesis (Fig. 5, A **and** B). The wtHtt was depleted by 50%, whereas mHtt was depleted by ∼80%. (Fig. 5C). Notably, wtHtt depletion did not alter the FMRP levels in control striatal cells, but mHtt depletion significantly diminished the FMRP levels in the HD-homo striatal cells (Fig. 5 D), which is consistent with mHtt role in the upregulation FMRP protein in HD (Fig. 4 D, E). We next wondered whether the upregulated global protein synthesis upon mHtt depletion was due to diminished FMRP in the HD-cells. To address this question, we generated a double knockout of mHtt/FMRP using CRISPR/Cas-9. We found increased puromycin incorporation in mHtt-alone depleted cells, which was like mHtt/FMRP double-depleted cells (Fig. 5, E **and** F). As shown before (Fig. 4 J **and** K), FMRP depletion alone did not significantly affect the protein synthesis in the HD-homo cells (Fig. 5, E **and** F).

Next, we hypothesized that enhanced protein synthesis in wtHtt/mHtt-depleted cells might be accompanied by altered ribosome profiles. Under basal conditions, we found no apparent differences in the ribosome profiles or any changes in the PS/MS ratio between CRISPR control cells and CRISPR wtHtt-depleted control cells (Fig. 5G; ***blue***). However, in a ribosome run-off assay in the presence of harringtonine, we found that wtHtt-depleted control cells showed a rapid increase in MS peaks compared with controls, and there was a significant decrease in the PS/MS ratio (Fig. 5G; ***red***). Similarly, the PS/MS ratio was not altered under basal condition in HD-homo cells (Fig. 5H; ***blue*)**, but in the presence of harringtonine, the PS/MS ratio showed significantly decreased in the mHtt-depleted HD-homo cells compared to CRISPR control HD-homo cells (Fig. 5H; ***red***). Unlike mHtt, the depletion of FMRP did not alter the PS/MS ratio in the HD-homo cells in basal or after harringtonine treatment (Fig. 5, I **and** J), which further suggests that FMRP does not regulate overall ribosome stalling in HD cells. These data indicate that depletion of wtHtt or mHtt enhances protein synthesis and increases the speed of ribosome movement. Therefore, normal Htt is a novel regulator of ribosome stalling, which is potentiated by mHtt, an effect that operates independent of FMRP.

### mHtt interacts directly with ribosomes *in vitro* and to an RNase-sensitive component of ribosomes in the brain

To investigate whether mHtt interacts with ribosomal proteins, we immunoprecipitated (IP) Htt from human HD (69Q^+/-^) and control (23Q^+/-^) fibroblasts using the validated MAB2166 antibody, which does not discriminate between wtHtt or mHtt, coupled with mass spectrometry (IP-LC-MS/MS). The cells were either starved for amino acids or starved and then stimulated with L-leucine (Leu), followed by the IP-LC-MS/MS. We found that Htt interacted with several 40S and 60S ribosomal proteins both in the starved and amino acid-stimulated conditions (Fig. 6A). STRING analysis found an enrichment of the biological process and components related to the translation and ribosome/RNA binding components (fig. S12, A **and** B). Full interactome data can be found in **Data file S12**. Next, using super-resolution STED microscopy, we found that around 25—35% of Htt clusters were in proximity (<300 nm) to the Rpl7 (Fig. 6B**, full image in** fig. S10B), with clear differences observable between the control and homo conditions when calculating the Manders’ Colocalization Coefficients (Fig. 6C). Next, by employing ribosome fractionations of striatal cells, we found both wtHtt and mHtt co-sediments with the 40S, 60S, and 80S (MS) subunits, and with PSs in the sucrose gradient in control and HD-het striatal cells (fig. S13). We validated that endogenous mHtt co-precipitated ribosomal proteins in a poly-Q dependent fashion, consistent with a previous study (46). Figure 6D shows Rps6 and Rpl7 strongly coprecipitated with mHtt in HD-homo cells compared with HD-het and control cells. As a positive control, we detected Caprin1, a previously known interactor of mHtt (47) (Fig. 6D). Next, we investigated whether Htt can directly interact with PSs. We incubated the GST-tagged exon 1 of recombinant Htt proteins (Exon 1 23Q or 51Q) (48) with PSs isolated from the striatal cells depleted of endogenous Htt (CRISPR-wtHtt) and re-ran the isolated PSs on a sucrose gradient (see flow chart fig. S14). We found that mHtt-51Q bound strongly to the extracted PSs, compared with wtHtt-23Q (Fig. 6E). This data indicates wtHtt and mHtt are associated with ribosomes in human HD fibroblasts and *in vitro* and mHtt show stronger affinity to ribosomes and ribosomal proteins in co-immunoprecipitation assays.

Next, we investigated how wtHtt and mHtt interacted with ribosomes in the HD mouse brain (zQ175DN) (45). As expected, the treatment of HD cortical brain lysate with EDTA, which disassembled the ribosomes (Fig. 6F), resulted in the relocation of the ribosomal proteins Rpl7, Rpl35, or FMRP to 60S and 80S (Fig. 6, G-H), confirming their association with PSs. Unlike these proteins, in EDTA, neither wtHtt nor mHtt (detected by the poly-Q antibody) resedimented to lower fractions, and there was a substantial amount of the EDTA-insensitive fractions of wtHtt/mHtt in the PS (Fig. 6, F-H), consistent with a previous study (46), suggesting that Htt might bind tightly to ribosome/membrane compartments that are known to be resistant to EDTA treatment (49). Treatment of a mouse HD cortical brain lysate with RNase, which promoted the degradation of PS-bound mRNA substrates (Fig. 6, F-H), resulted in the accumulation of Rpl35, Rpl7 as expected in the lower density fractions. mHtt, somewhat resembling FMRP, also accumulated in the lower density fractions (Fig. 6G**, arrow**), but a substantial portion of wtHtt remained in the higher density polysome fractions (Fig. 6, G **and see graphs in H**). These data indicate that wtHtt and mHtt bind differentially to RNA-protein components of ribosomes, and that mHtt interaction, but not wtHtt with ribosomes, is RNase-sensitive (**Compare insets g1 and g2**). Collectively, these data indicate that enhanced inhibition of protein synthesis and ribosome stalling in HD might stem from the much stronger interaction of mHtt to the RNase-sensitive components of ribosomes.

## DISCUSSION

Based on the data presented above, our model suggests that wtHtt plays a major role in the physiological inhibition of protein synthesis by regulating the speed of ribosome translocation on mRNAs. However, this function is remarkably enhanced by glutamine expansion, which promote slow and stalled ribosomes, resulting in a “toxic gain of inhibition” by mHtt (Fig. 6I). Also, our Ribo-Seq data revealed the positions of the ribosomes on mRNA; therefore, for the first time, we have demonstrated unique ribosome pauses in HD cells at a single-nucleotide resolution on selected mRNA transcripts. It is well known that ribosomes stall at discrete sites on mRNA for reasons that are less well understood (50, 51). In HD cells, we found ribosome pauses on several mRNA transcripts at both the single-codon and the gene level; the HD-based ribosome pauses of the mRNA targets are specific to HD. We arrived at this notion because the ribosome pausing was not affected by already known pause targets, such as Srpr or sec11a, which showed predicted pauses (high RPF density) in their ORF due to co-translation translocation events at exons 3 and 4 (52), respectively, both in controls and the HD cells (fig. S15, A **and** B). Similarly, we found a discrete distribution of RPF on apoB mRNA (fig. S15C), analogous to 23 translocational pauses previously demonstrated in an in vitro translation system (53). Therefore, RPF in our Ribo-Seq showed predicted and HD-specific ribosomal pauses.

What are the mechanisms by which mHtt promotes ribosome stalling? Many factors might be involved (including secondary RNA structures, 3’ UTR, or codon usage), or the availability of charged tRNAs, ribosomal binding proteins, and/or nascent polypeptide chains may contribute to ribosome pauses (54). Answering key questions, such as identifying signature mRNA motifs, the nature of protein partners, or mRNAs on the PSs that wtHtt or mHtt interact with, will help identify the mechanisms. At least we know that FMRP, a known regulator of ribosome stalling (55), appears not to be involved in mHtt-mediated ribosome stalling. Its subtle effect in *in vivo* suggests that mHtt and FMRP might alter a selective set of mRNA transcripts, for example, synapse-enriched targets (56), to regulate motor skill learning in HD (Fig. 4Q). It is entirely possible that mHtt and FMRP stall ribosomes using different mechanisms, or they might regulate translation independent functions in the brain (57). Consistent with this notion, a recent study showed wtHtt mediates FMRP-dependent regulation of mitochondrial fusion and dendritic maturation (43). Collectively, the current study’s results revealed that mHtt promotes ribosome stalling and affects a diverse set of mRNAs transcripts, which might lead to the progressive and widespread development of HD-related behavioral and pathological symptoms. Therefore, the identification of ribosome stalling mechanisms and the elucidation of the role of paused mRNA transcripts in HD cells may help identify novel therapeutic targets to prevent or slow the progress of HD.

## Acknowledgments

We would like to thank Melissa Benilous for administrative help, and members of the lab for continuous support and collaborative atmosphere. We like to thank members at the Scripps proteomics and genomic core for their help and expertise. This research was supported by grant awards from NIH/NINDS R01-NS087019-01A1, NIH/NINDS R01-NS094577-01A1, and the Cure for Huntington Disease Research Foundation (CHDI).

## Author contributions

S.S made the initial observations and further conceptualized and co-designed the project with M.E. M.E carried out all the polysome profiles, and related biochemical experiments, and together with P. K generated cDNA library. E. R and J. B carried out bioinformatics analysis. A.M. generated the data for the triplet periodicity, metagene, pause analysis, center of ribosome density and differential gene expression analysis. N.S and S.R.R carried out IP proteomics, N. G carried out validation blot experiments. N. U carried out STED imaging and co-localization analysis. K.F supported in profiling and blotting. M.E and S.S analyzed the data. S.S wrote the paper with input from co-authors.

## Data and materials availability

All the data is available in the main text or the supplementary materials.

## Supplementary Materials

### Supplementary Text

A selected profile from other mRNAs shows an intriguing distribution of ribosomes across the mRNA. For example, the autism susceptibility candidate 2 (Auts2), which is associated with autism and mental retardation (58), showed ∼one fold RPF increase concentrated on exon 2 in HD-Het compared to control cells (**fig. S5A, arrow**). The adrenergic receptor, alpha 2a, contains one long exon, which regulates dopamine/DARPP-32 signaling in the striatum (59), showed a robust RPF upstream of annotated exon 1 start codon in HD-homo and HD-het cells (fig. S5B, arrow). The CASP8 and FADD-Like Apoptosis Regulator (Cflar), a regulator of apoptosis upregulated upon inhibition of protein translation (60), showed RPF on exon 4 in HD-homo and HD-het cells compared to control cells (**Fig. S5C, arrow)**. Ddit4l, DNA Damage Inducible Transcript 4, a. k. a REDD1), an inhibitor of mTORC1 signaling (61), showed an ∼one-fold higher RPF in HD-Homo and compared to HD-het, with an enrichment of RPF on exon 2 and 3 (arrow), but a very few read in control (**fig. S5D**). Finally, the NEDD4-like, E3 ubiquitin-protein ligase, Wwp1, which is involved in the trafficking of proteins to proteasomes or lysosomes (62), consists of 25 exons, but displayed a signature concentration of RPF in exon 1 in all groups (**fig. S5E**). A full UCSC Genome Browser link for the global ribosome footprints for HD and control cells can be found in this link: Https://genome.ucsc.edu/cgi-bin/hgTracks?hubUrl=Https://de.cyverse.org/anon-files/iplant/home/rmi2lab/Hub_Collaborations/Srini/hub.txt&genome=mm10

### Materials and Methods

#### Cell culture

Mouse striatal cells (ST*Hdh*) expressing knock-in wild-type Htt^exon1^ with 7 Glu repeats (control; ST*Hdh^Q7/Q7^*) or expressing knock-in mutant human Htt^exon1^ with 111 Glu repeats (HD-het; ST*Hdh^Q7/Q111^*, and HD-homo; ST*Hdh^Q111/Q111^*) (25) were purchased from Coriell Institute for Medical Research (Camden, New Jersey, USA) and cultured in 10% FBS, DMEM, high glucose, 5% CO2, at 33°C, as described before (24).

#### RNase foot printing

Global RNase foot-printings were performed during three independent rounds of cell cultures (n=3). For each round of global foot printing, mouse immortalized striatal cells (i.e. control, HD-het, and HD-homo cells) were plated in 15 cm dishes at a confluency of 70%. The following day the mediums were changed and after 2 hours the cells were incubated with cycloheximide (CHX, 100 μg/ml) for 10 min. The cell then scraped and washed with cold PBS (containing 100 μg/ml CHX) twice. During the second wash 5% of cells were transferred to new tubes and were lyzed by adding 700 µl of QIAzol lysis reagent. Total RNAs of these samples were isolated using miRNeasy Mini Kit (Qiagen) for total mRNA sequencing. After the second wash, the rest of the cells were lyzed in a lysis buffer containing 20 mM HEPES pH 7.3, 150 mM KCl, 10 mM MgCl_2_, 2mM DTT, 100ug/ml Cyclohexamide, 0.5% v/v Triton X-100, 20 U/ml RNasin and EDTA free protease inhibitor cocktail (Roche). The cell lysates were passed 20 times through a 26G needle and incubated on ice for 15 minutes, then centrifuged at 21000 rpm for 15 minutes. Supernatants were transferred to new tubes. Equal total RNA amount of each sample was used for global RNase foot printing as follow; for each A260 absorbance unit of the lysates 60 units of RNaseT1 (ThermoFisher Scientific) and 0.6 µl of RNaseA (Ambion) were added and the samples were incubated at 25C for 30 min. RNase treated samples were immediately loaded on 10-50% sucrose gradients and centrifuged at 40000 RPM (SW41Ti rotor) at 4°C for 2 hours. Gradients were fractionated using a gradient fractionator and UA-6 detector (ISCO/BRANDEL). Fractions containing 80S peaks of each sample were collected and their RNAs were isolated using a miRNeasy Mini Kit (Qiagen).

#### List of antibodies

Chemicals and reagents were mainly purchased from Sigma. Htt protein was produced as described before (63). Huntingtin (MAB2166) and puromycin (MABE343) antibodies were obtained from, Millipore-Sigma. Anti-polyglutamine antibody (P1874) was from Sigma. Actin (sc47778), Plectin (sc33549), CNTF (sc25286), Clic4 (sc271863) and GST-horseradish peroxidase (HRP, sc138 HRP) antibodies were from Santa Cruz Biotechnology. RPL7 IHC Antibody (IHC-00455), RPL35A/Ribosomal Protein L35a Antibody (A305-106A), Caprin1 (A303-881A) from Bethyl Laboratories; Lrrc17 (20918-1-AP), Phf11d (10898-1-AP), Acta2 (14395-1-AP), Mfsd10 (11518-1-AP) were from Proteintech. mTOR (2972), FMRP (4317), S6 (2217), eIF2α (5324), P-eIF2α (3398) and cGAS (31659) were from Cell Signaling Technology. HRP-conjugated secondary antibodies were from Jackson ImmunoResearch Inc. Secondary antibody for STED microscopy were Anti-mouse STAR 635p, Anti-rabbit Alexa 594.

#### Generation of cDNA libraries from ribosome protected mRNAs

The following procedure were performed for all the RNA samples simultaneously. 20 µg of each sample was run on a 15% TBE-Urea gel (Novex) along with 26 and 32 nt RNA markers. The gel containing each sample was excised between two markers. RNAs were extracted from gel pieces by incubating gel slurries with nuclease-free water overnight at 4C and precipitated using RNase-free isopropanol and then eluted in nuclease-free water. T4 Polynucleotide Kinase (NEB) was used to catalyze the addition of 5’ monophosphate and removal of the 3’ phosphate in the RNA fragments to leave a 3’ hydroxyl terminal needed for adapter ligation. RNA was purified using the Zymo clean and conc-5 kit (Zymo Research, Cat. # R1013). Ribosomal RNA was depleted from the samples using TruSeq total RNA rRNA-depletion protocol (Illumina, Cat. #RS-122-2201) and then RNA samples were purified using Agencourt RNAClean XP beads (Beckman Coulter).

#### Generation of cDNA libraries and sequencing

NEXTflex small RNA-seq Kit v3 (Perkin Elmer) was used to ligate 5’ and 3’ adapters to purified RPF fragments, which then were reverse transcribed and amplified (14 cycles) to generate cDNA libraries. Libraries were cleaned up using NEXTflex Cleanup beads, pooled and sequenced in the NextSeq 500 (V2) using single-end 50bp chemistry at the Scripps Genomic Core, at Florida, USA.

#### Generation of mRNA-seq libraries

NEBNext Ultra II Directional kit (NEB, Cat. # E776)) with the NEBNext poly(A) mRNA Magnetic isolation module (NEB, Cat. # E7490) was used generate mRNA-seq libraries. Briefly, 400ng of high-quality total RNA was used to purify poly(A) mRNA, fragmented, reverse-transcribed with random primers, adapter ligated, and amplified according to manufacturer recommendations. The final libraries were validated on the bioanalyzer, pooled, and sequenced on the NextSeq 500 using paired-end 40bp chemistry.

#### Ribo-Seq, RNA-seq quality control and mapping the reads to UCSC browser

RNAseq reads were trimmed using Cutadapt (64) with the following parameters : -a AGATCGGAAGAGCACACGTCTGAACTCCAGTCA -A AGATCGGAAGAGCGTCGTGTAGGGAAAGAGTGT --minimum-length=15 –pair-filter=any. For Riboseq reads, 3’ adapters were trimmed using Cutadapt with the following parameters : -a TGGAATTCTCGGGTGCCAAGG --minimum-length 23. The reads were further trimmed using Cutadapt to remove 4 bases from either side of each read accordingly to the NEXTflex™ Small RNA Trimming Instructions (cutadapt -u 4 -u −4). Fastq files were checked for quality control with FastQC. Both RNAseq and Riboseq reads were next mapped to a library of mouse rRNA and tRNA sequences using Bowtie v1.1.2. Any reads mapping to these abundant contaminants were filtered out. Remaining reads were then aligned to the mouse transcriptome with RSEM v1.3.0 (65) using the GRCm38.p5 genome annotation and the comprehensive gene annotation from Gencode (M16 release) as transcriptome reference. Reads with a mapping quality <5 were discarded. Cleaned bam files were converted to bigWig files with Bedtools (66) for visualisation using the UCSC Genome Browser. For the euclidian distance analyses, gene expression was quantified with RSEM and comparison plots were generated in R using DESeq2 (67) and ggplot2 packages.

#### Ribosome occupancy (Anato2Seq) analysis

The raw Ribo-seq reads were clipped of adapter sequence (TGGAATTCTCGGGTGCCAAGG) using Cutadapt (version 1.18) (64) with the following command: cutadapt -f fastq -a CTGTAGGCACCATCAAT --minimum-length=23 <input>.fastq -o <output>.fastq. A 4bp secondary trim from either end of the reads was performed also using Cutadapt. Mouse rRNA sequences were retrieved from NCBI (cite PMID: 29140470) with the following accessions: NR_003279, NR_003278, NR_003280, NR_030686. Ribo-seq and RNA-seq reads aligning to these sequences were removed using bowtie (version 1.0.1) (cite PMID 19261174) with the following command: bowtie - v 3 --norc <path_to_rRNA_indices> -q <input>.fastq --un <output>.fastq. The remaining reads were then mapped using bowtie to the RefSeq (cite PMID: 26553804) mouse transcriptome downloaded from ftp://ftp.ncbi.nlm.nih.gov/refseq/M_musculus/mRNA_Prot/ on 9th October 2018. The following command was used: bowtie -a -m 100 -l 25 -n 2 -S --norc <path_to_transcriptome_indices> -q <input>.fastq <output>.sam.

The sam alignment files were parsed with an in-house python script to count the number of reads aligned to each gene using an exon union approach (cite PMID: 25943107). The Ribo-Seq reads were assigned to mRNA coordinates using an offset of 14 nucleotides downstream of the 5’ end of the reads. The counts included uniquely mapped reads i.e. reads that mapped to one location only in the mouse transcriptome plus reads that mapped up to 3 locations in the transcriptome which were weighted by the number of their mapped locations (up to 3).

For differential expression analysis using anota2seq (cite https://www.biorxiv.org/content/10.1101/106922v4), the number of reads aligning to annotated CDS regions was used for Ribo-seq while for RNA-seq the number of reads aligning to the entire transcript was used. These counts were input to anota2seq (version 1.5.2) with the parameters dataType = “RNAseq”, filterZeroGenes = FALSE, normalize = TRUE, transformation = “TMM-log2”.

#### Ribosome pause (PausePred) analysis

The command line version of the PausePred software (PMID: 30049792) was run with the following parameters for each replicate (control, HD-het and HD-homo): fold change : 5; window size: 1000 nucleotides; read lengths: 26-32 nucleotides; window coverage: 5. Individual offset values were assigned according to metagene analysis for each read length (26-32 nucleotides) accounting for mismatches at the 5’ ends of the reads.

For genes with detected pauses, the center of ribosome density (cite PMID: 25621764) was determined using an in-house python script.

#### Deletion of *Hdh* (Htt) and *Fmr1* (FMRP) genes using CRISPR/Cas9

GFP expressing CRISPR/Cas9 knockout plasmids for mouse *Htt* and *Fmr1* were purchased from Santa Cruz Biotechnology (#sc-420825 and #sc-420392 respectively). Mouse striatal cells were transfected with knockout plasmids by PolyFect (Qiagen) following recommendations by the company. The transfected cells were harvested 48 hrs later, washed and resuspended in the Sorting Buffer buffer (containing 1x Phosphate Buffered Saline (Ca/Mg++ free), 2.5 mM EDTA, 25 mM HEPES pH 7.0, 1% Fetal Bovine Serum and 10 unit/ml DNase1). The GFP expressing cells were sorted using a BD biosciences Aria FACs machine. The sorted cells were plated and maintained in DMEM containing 10% FBS at 33°C.

#### Immunostaining of the cells and STED image acquisition

Mouse striatal cells were plated in 12 well plates containing glass coverslips. The day after the cells were washed with PBS and fixed using 4% PFA in PBS, then permeabilized in 0.1% Triton X-100 in PBS and blocked using 5% normal donkey serum/ 1% BSA/ 0.1% Tween in PBS. The cells were incubated with primary antibodies (FMRP 1:50; CST #4317; Rpl7 1:50; Bethyl lab # IHC-00455) in blocking buffer at 4 degrees overnight. The day after, the cells were washed and incubated with secondary antibodies and DAPI in 1% BSA/ 0.1% Tween in PBS for 1 hr at the room temperature. The stimulated emission depletion (STED) microscopy imaging was performed using a multicolor Expert-Line STED nanoscope (Abberior Instruments GmbH), using the 775-nm pulsed STED laser in combination with the 561-nm and 640-nm pulsed excitation lasers, as well as the 405-nm excitation for diffraction-limited imaging of DAPI. All images were recorded simultaneously with diffraction-limited (i.e., confocal) and with 2D-STED enhanced resolution, recording each color channel and resolution in a line-interleaved manner. Pixel sizes (x,y) were 20-nm x 20-nm for both STED and confocal images, with typical pixel dwell times of 10 µs for confocal and 30—45 µs for STED images. The images were recorded with a 100× oil immersion objective lens (NA 1.4), using the QUAD beam scanner, utilizing the Imspector software package (Max-Planck Innovation).

#### Image analysis

For the colocalization analysis, raw STED images from Imspector were imported into FIJI and were processed as follows. First, raw images were gently smoothed with a 1-pixel Gaussian filter. Next, an appropriate background level was determined individually for each image, striking a careful balance between being able to distinguish individual clusters in proximity without losing any of the dimmer features. This background value was then subtracted from the images. For some parts of the analysis, the images were then binarized. Regions coinciding with the cell nucleus were excluded from the colocalization analysis, as were smaller regions coinciding with any obvious staining artifacts. The colocalization analysis itself was performed using the ImageJ plugin JACoP, utilizing both the pixel-based Manders coefficient analysis and the object-based methods.

#### Ribosome run off assay

Mouse striatal cells were plated at 60-70 % confluency, on next day were incubated with harringtonine (2 μg/ml final concentration) for different time points at 37°C. The cells were immediately incubated with CHX (100μg/ml) for min and then scraped. Polysome profiles for each sample was prepared as described before.

#### cDNA preparation and Real-time PCR

A sucrose density gradient centrifugation was carried out using control and HD-homo cells and ribosomes fractions were collected. RNA was extracted from the fractionated samples following lysis in Trizol reagent. 250 ng RNA was used to prepare cDNA using Takara primescript^tm^ kit (Cat no. 6110A) using random hexamers. The qRT-PCR of genes was performed with SYBR green (Takara RR420A) reagents. Primers for all the genes were designed based on sequences available from the Harvard qPCR primer bank. The primer sequences are as follows: Gapdh mouse (Forward primer) 5’ primer AGGTCGGTGTGAACGGATTTG (Reverse primer); 3’ primer TGTAGACCATGTAGTTGAGGTCA: Fmr1 mouse (Forward primer) 5’ primer CCAATGGCGCTTTCTACAAG; (Reverse primer) 3’ primer TCTGTCTCTCTGGTTGCCAGT: Rps27 mouse (Forward primer) 5’ primer ACGACCTCCCTACGAGAACA; (Reverse primer) 3’ primer ATAGCATCCTGGGCATTTCA.

#### Protein precipitation from sucrose fractions

Total Proteins were precipitated from monosome/polysome sucrose fractions using methanol/chloroform. Protein pellets were resuspended in a buffer containing 10mM Tris-HCl pH 8 and 0.1% SDS.

#### Puromycin metabolic labeling and immunoblotting

Mouse striatal cells were plated at a confluency about at 60-70%. The day after the cells were incubated with puromycin (20 μM final concentration) for 5 min as described (26). Then cells were rinsed with cold PBS and immediately were lysed in RIPA buffer containing protease inhibitors. Equal proteins were used to run western blotting experiments.

#### Immunoprecipitation striatal cells

Control, HD-Het and HD-Homo striatal cells (2 x 10^6^) were plated in 10-cm dishes and next day were lysed in immunoprecipitation (IP) buffer [15 mM HEPES (pH 7.3), 7.5 mM MgCl_2_, 100 mM KCl, 1.0% Triton X-100, 1 mM dithiothreitol (DTT), EDTA-free protease inhibitor cocktail (Roche), RNasin (40U/μl, Promega). The lysates were run several times through a 26-gauge needle in IP buffer and incubated on ice for 15 min and centrifuged 11,000 rpm for 15 min. Protein estimation in the lysate supernatant was done using a bicinchoninic acid (BCA) method, a concentration (1 mg/ml) of protein lysates was precleared with 40 μl of protein A/G beads for 1 hour, supernatant was incubated for 1 hour at 4°C in Htt IgG (MAB2166) or control IgG, and then 60 μl protein A/G beads were added and incubated overnight at 4°C. After 12 hours, the beads were washed five times with IP buffer (without RNasin/protease inhibitor), and the protein samples were eluted with 30 μl of 2x lithium dodecyl sulfate (LDS) containing +1.5% β-mercaptoethanol, separated on NuPAGE 4-12% Bis-Tris gel (Thermo-Fisher Scientific), transferred to polyvinylidene difluoride membranes, and probed with the indicated antibodies. HRP-conjugated secondary antibodies (Jackson ImmunoResearch Inc.) were probed to detect bound primary IgG with a chemiluminescence imager (Alpha Innotech) using enhanced chemiluminescence from WesternBright Quantum (Advansta).

#### Immunoprecipitation in human fibroblasts and LC-MS/MS

HD patient-derived fibroblast cell lines GM04281 (wild type HTT allele/17 CAG repeats, mutant HTT allele/69 CAG repeats) and normal human fibroblast cell line GM07492 were obtained from the Coriell Institute (Camden, NJ). Cells were maintained at 37°C and 5% CO2 in Dulbecco’s modified Eagle’s medium, high glucose, GlutaMAX supplement (DMEM) (Thermo Fisher Scientific) supplemented with 10% fetal bovine serum (Thermo Fisher Scientific), 1% penicillin-streptomycin and 1% MEM nonessential amino acids (Thermo Fisher Scientific). Human fibroblasts were plated in 10-cm dishes and next day the medium was changed to Krebs buffer medium [20 mM HEPES pH 7.4, glucose (4.5 g/liter), 118 mM NaCl, 4.6 mM KCl, 1 mM MgCl_2_.6H_2_O, 12 mM NaHCO_3_, 0.5 mM CaCl_2_, 0.2% (w/v) bovine serum albumin (BSA)] devoid of serum and amino acids for 1 hour to simulate full starvation conditions. For the stimulation conditions, cells were stimulated for 15 min with 3mM leucine. Cells were lysed and proceeded for immunoprecipitation as mentioned above for the striatal cells. After running the IP samples in electrophoresis, the samples were subjected to IP–LC-MS/MS as described previously (68)) for the analysis of HTT interactors.

#### Immunoprecipitation human HD brain

Human brain tissue (Caudate nucleus) samples of grade 1 HD-affected patient (HSB # 3358, 2706, 3744), grade 2 HD-affected patient (HSB # 2858, 3432, 3635, 3872, 4072), grade 3 HD-affected patient (HSB # 4344, 2869, 2972, 4518), grade 4 HD-affected patient (HSB # 5078, 2903) and normal donor controls (HSB # 4615, 4823, 5293, 4340, 4135) were obtained from the Human Brain and Spinal Fluid Resource Center VA West Los Angeles Healthcare Center, Los Angeles, CA. Human tissue was homogenized in binding/lysis buffer [50 mM tris (pH 7.4), 150 mM NaCl, 10% glycerol, and 1.0% Triton X-100] with protease and phosphatase inhibitors, followed by a brief sonication for 6 s at 20% amplitude. The lysates were subjected to Htt immunoprecipitation as mentioned above for the striatal cells.

#### Ribosome isolation and in vitro binding assay

For each assay four 15 cm plates of mouse striatal cells were used. Briefly the cells were incubated with 100 μg/ml CHX for 10 min, then harvested, spin down and washed once with cold PBS containing CHX. The cells were lyzed in the lysis buffer containing 20 mM HEPES pH 7.3, 150 mM KCl, 10 mM MgCl_2_, 2 mM DTT, 100 μg/ml CHX, 0.5% v/v Triton X-100, 20 U/ml RNasin and EDTA free protease inhibitor cocktail (Roche). The cell lysates were loaded on 10-50% sucrose gradients and centrifuged at 40000 RPM (SW41Ti rotor) at 4°C for 2 hours. The fractions containing monosome and polysomes were collected and transferred to a new 50mL falcon tube and were diluted using isolation buffer containing 20 mM HEPES pH 7.3, 150 mM KCl, 10 mM MgCl, 2 mM DTT, 100 μg/ml cyclohexamide (at least 1 in 3 for the monosome fractions and 1 in 5 for the polysome fractions). The diluted fractions were put on top of 1M sucrose cushion (made in isolation buffer) and centrifuged at 31500 RPM (SW32Ti rotor) at 4°C overnight. The pellets were rinsed gently with isolation buffer, then incubated with 50 μl of ribosome isolation buffer and stored on ice for 1 hr to allow the resuspension of the isolated ribosomes. Isolated ribosomes (50 nM final concentration) were incubated with recombinant proteins (500 nM final concentration) in isolation buffer for 10 min at room temperature. The samples were loaded on top of 10-50% sucrose gradients and centrifuged at 40000 RPM (SW41Ti rotor) at 4°C for 2 hours. The fractions containing monosome were collected. Protein were precipitated and used to run western blotting assays.

#### Animals and behaviors

The Q175DN HD mouse (B6J.zQ175DN KI #029928) and Fmr1^-/-^ (B6.129P2-Fmr1^tm1Cgr^/J # 003025) were purchased from the Jackson Laboratory and maintained and bred in Animal Research Core, Scripps Research, Jupiter, FL following the institute regulations. To generate Q175DN-Het;Fmr1^+/-^ mice, male Q175DN-Het and female Fmr1^-/-^ mice were crossed. The behavioral testing included body weight measurement, open field rotarod and inverted grip strength. Around 10 months old, Wt, Q175DN-Het, Fmr1^+/-^, or, zQ175DN-Het; Fmr1^+/-^ were subjected to rotarod, open field or inverted grip. Animals ability to stay on an accelerating rotarod (4-40 rpm) was examined for four consecutive days (maximum 300 s, 3 trials per session). Open field was conducted as described in our previous study (68). The inverted grip test was performed based on the Treat-NMD guidelines. Briefly, one mouse was placed on a metal grid. Then the grid was inverted gently and the latency to fall time was recorded (maximum 360 sec).

#### Statistical analysis

Data were expressed as mean ± SEM as indicated. All experiments were performed at least in biological triplicate and repeated at least twice. Statistical analysis was performed with a Student’s t-test or repeated measure two-way ANOVA followed by post-hoc Bonferroni multiple comparison test (Graphpad Prism7) as indicated in the figure legends.

**Figure S1-S15**

**External Databases S1-S12**

**fig S1.**
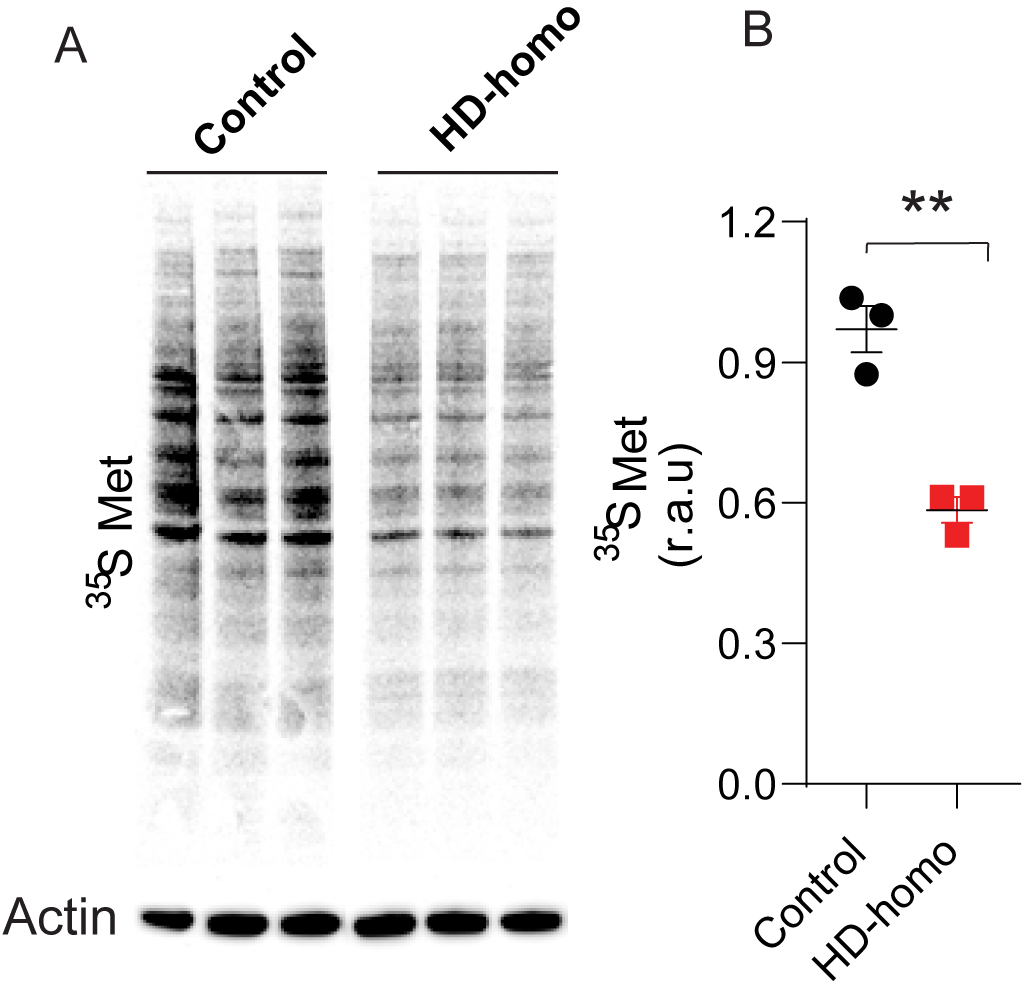
A. Representative autoradiograph of ^35^S Met in control and HD-homo striatal cells, and its quantification in B. The relative arbitrary unit, r. a. u.** P< 0.01 Student’ *t*-test.

**fig S2.**
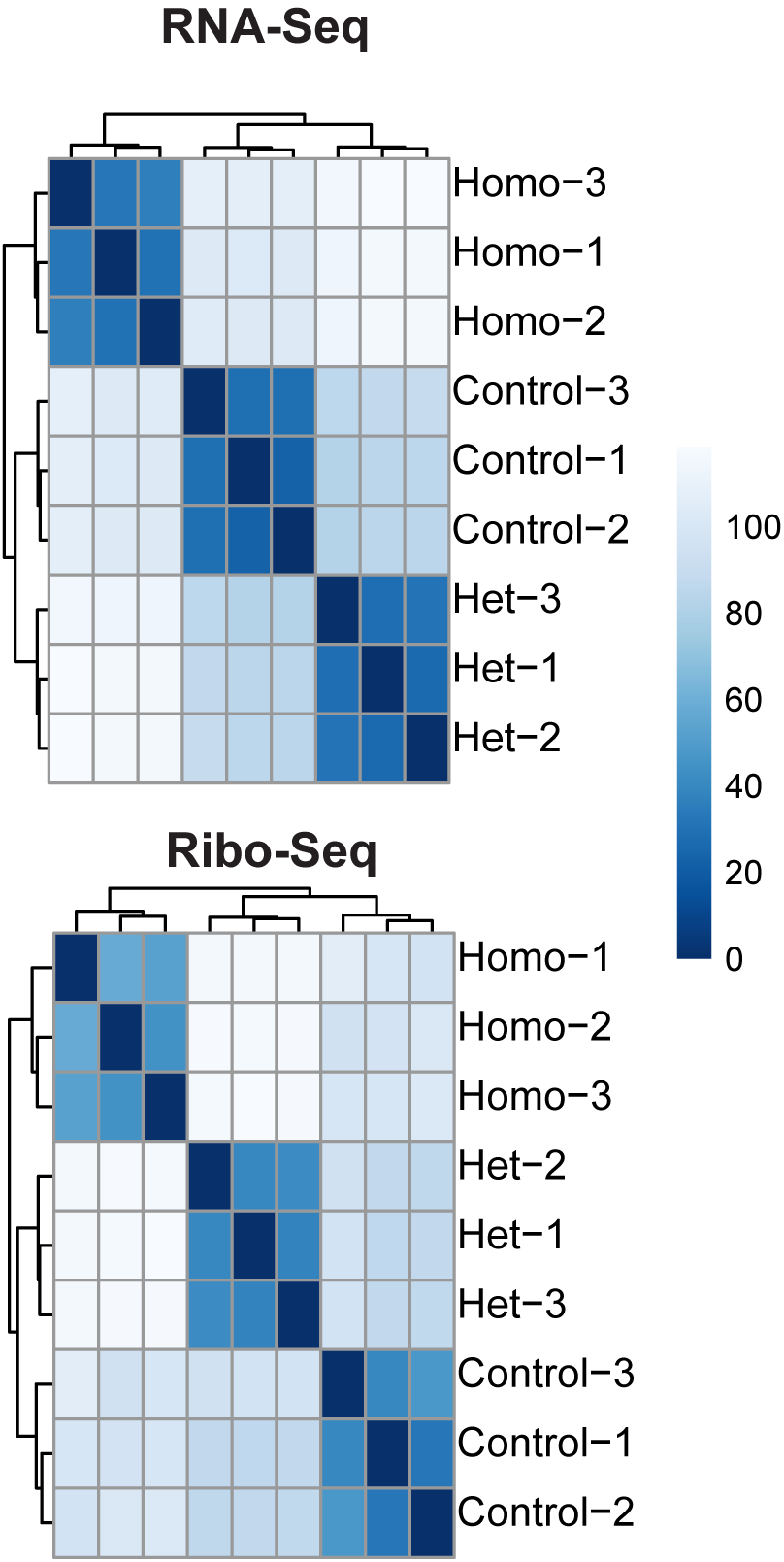
Euclidian distance analyses between the biological replicates.

**fig S3.**
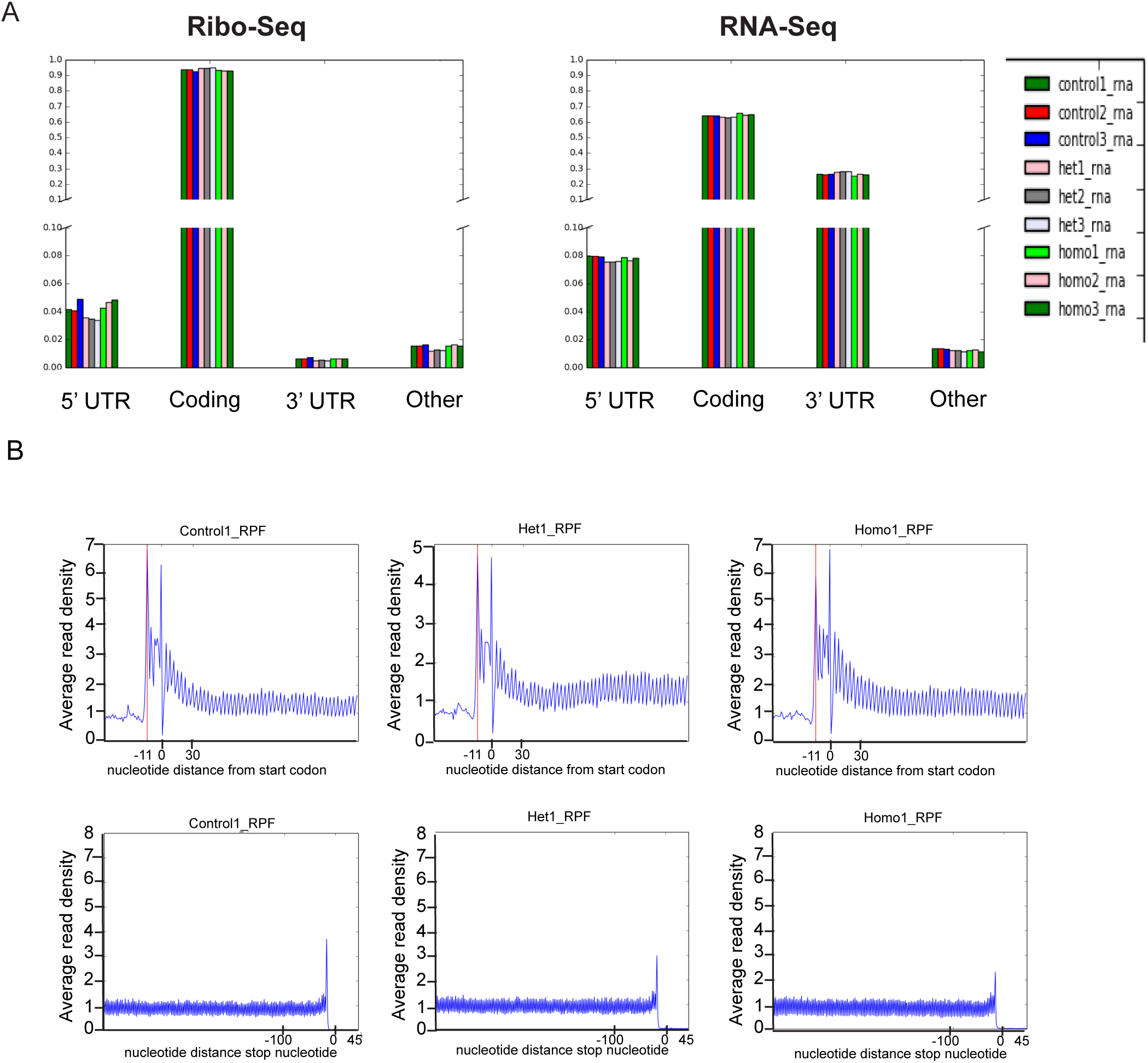
A. Ribo-Seq and RNA-Seq showing most reads are in coding region, B. Ribo-Seq showing ribosome occupancy at the start and stop codon.

**fig S4.**
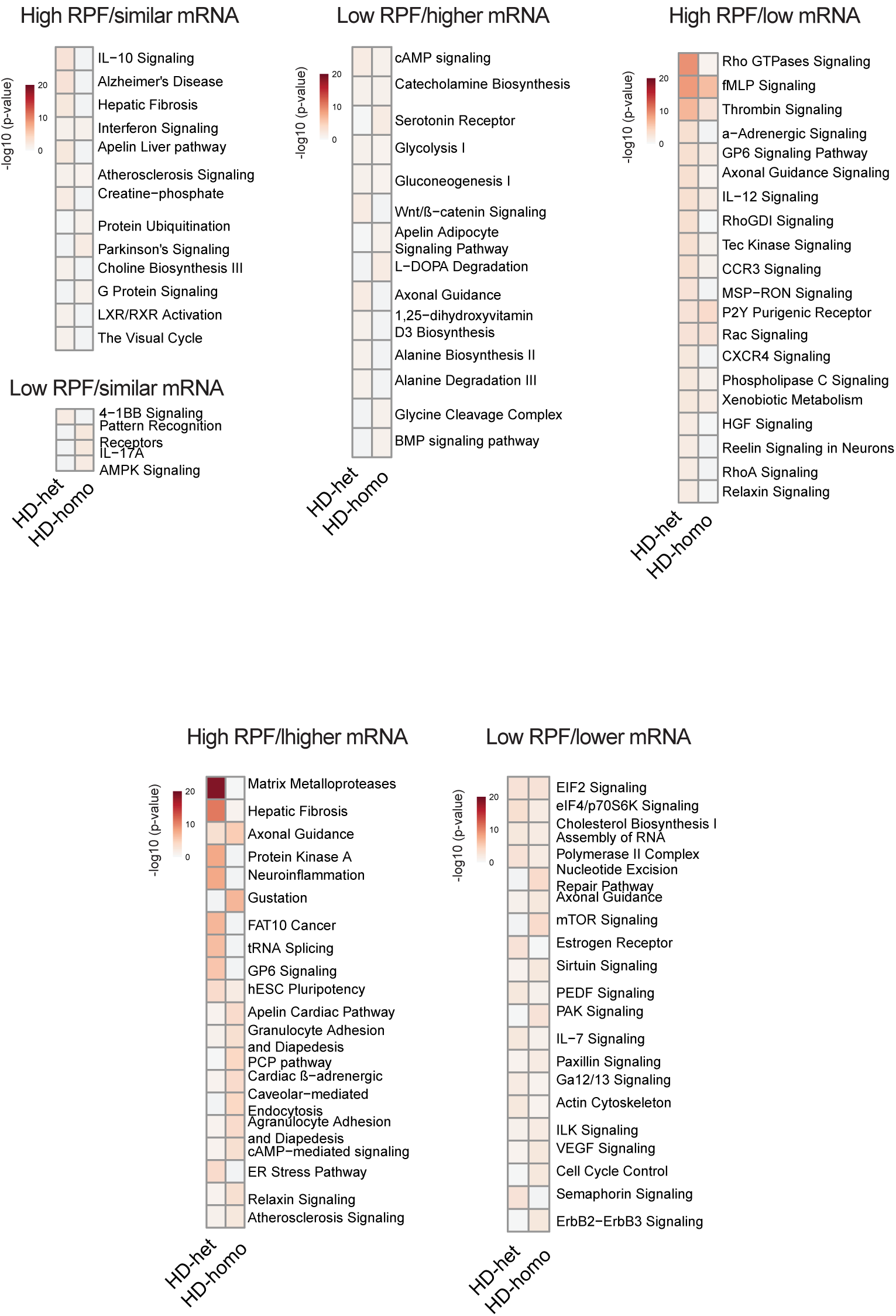
Gene-ontology (GO), using Ingenuity Pathway Analysis (IPA) in each category, showed genes (Data files S1-S4) belonged to diverse signaling pathways as indicated.

**fig S5.**
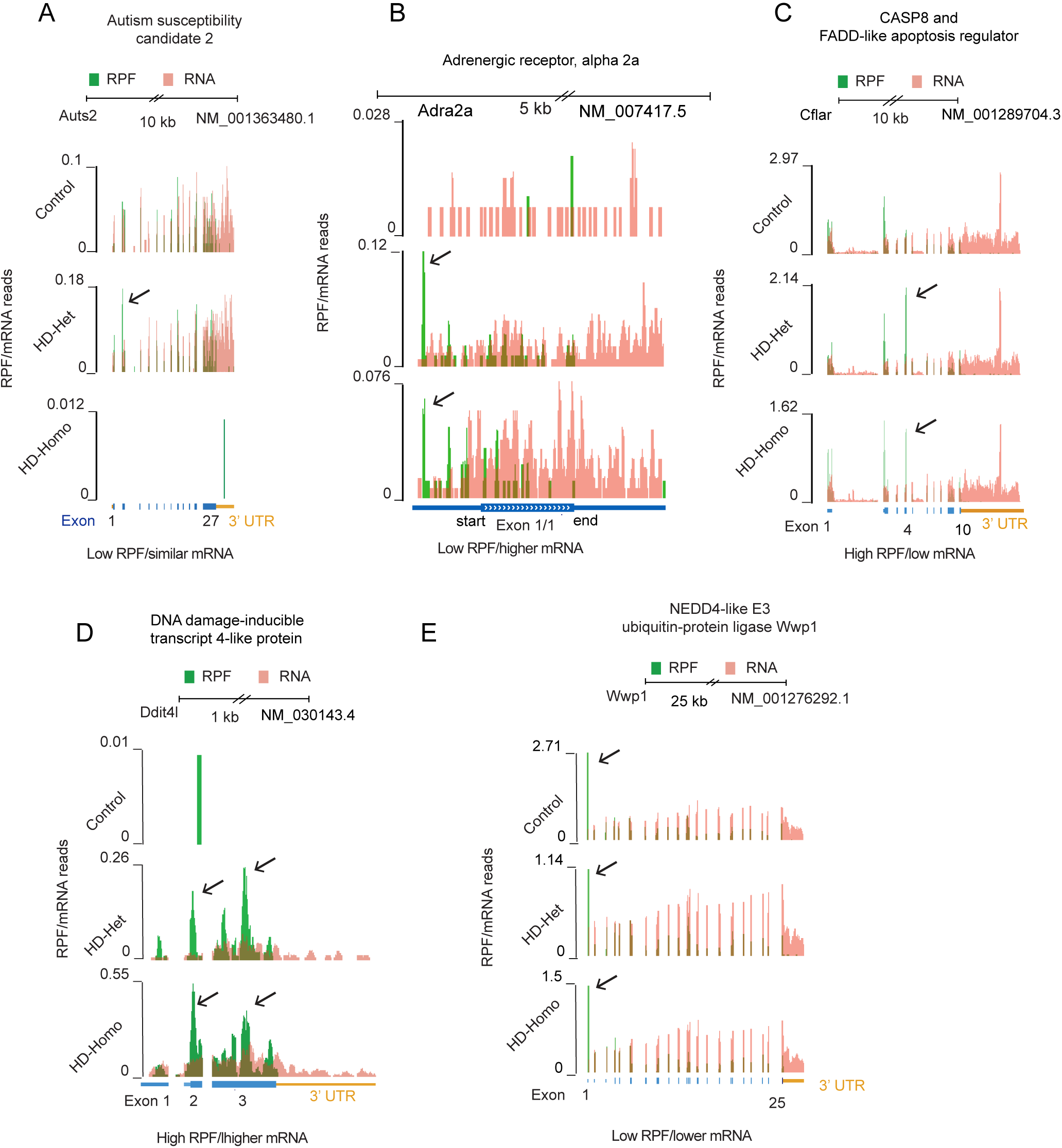
Overlay of RPF/mRNA profiles indicated genes from UCSC browser. A, the autism susceptibility candidate (*Auts2*). B, the Plecktrin homology domain containing family A member 4 (*Plekha 4*). C,the CASP8 and FADD-Like Apoptosis Regulator (*Cflar*), D, the DNA Damage Inducible Transcript 4, *Ddit4l* (a. k. a. *Redd1*). E, the NEDD4-like, E3 ubiquitin-protein ligase, *Wwp1*. Arrow indicates high ribosome occupancy.

**fig S6.**
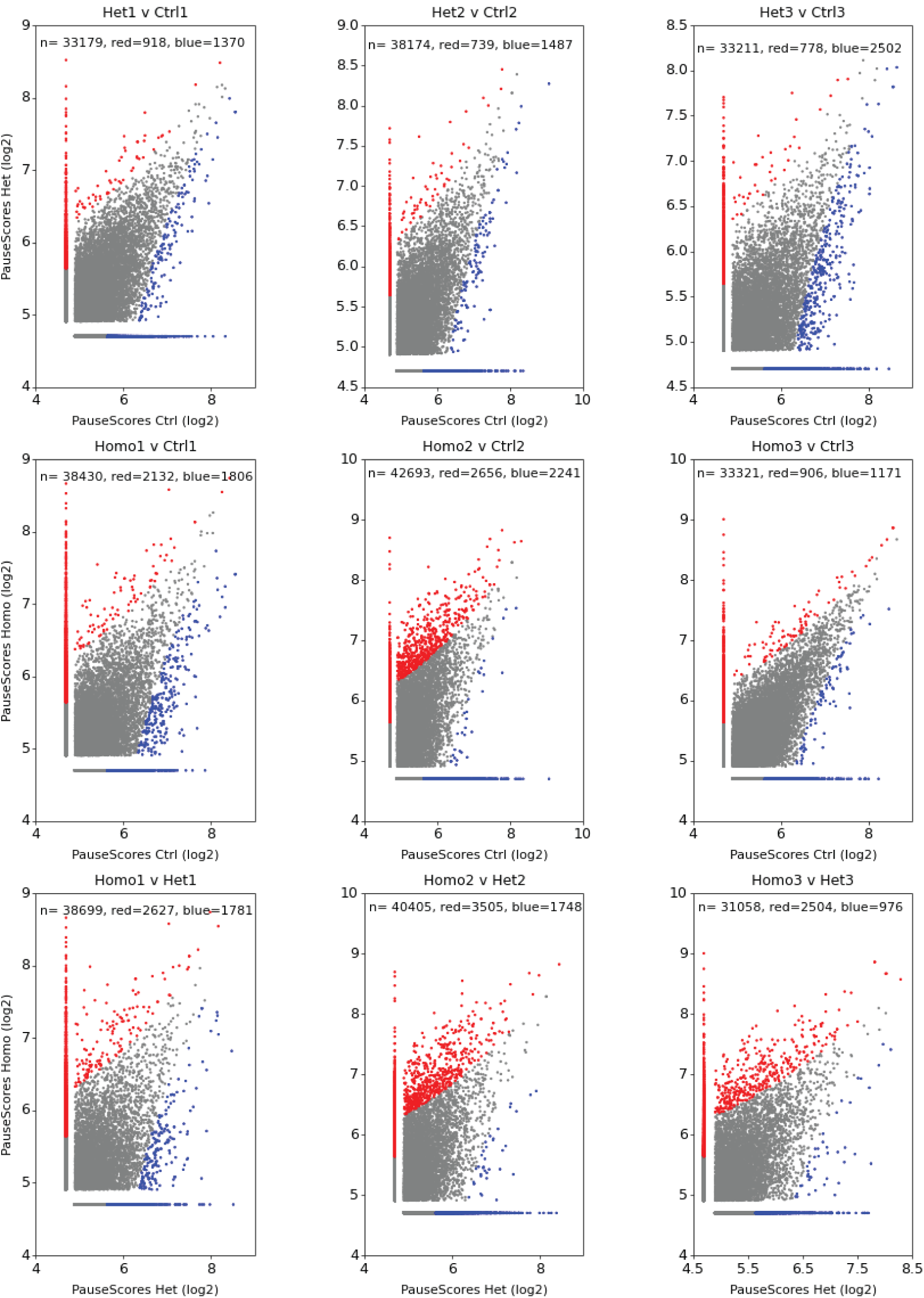
Pause difference within the control and HD replicates show single codon pause score (fold change) of 30 for the reads of length 26-32 nucleotides with the base pair coverage requirement of 20% within a window of 1000bp. When a pause is detected in e.g in het1 but not in ctrl1, it is represented in the grey/blue bar along the horizontal axis. The vertical grey/red bar represents those cases where the pause is detected in het1 but not in ctr1. The region between these bars represent pauses that were detected in both het1 and ctrl1. The red cases represent pauses in Het1 that have a pause score (fold change) of >=30 compared to the same location in ctrl1. Likewise, the blue cases represent pauses in ctrl1 that have a pause score (fold change) of >=30 compared the same location in het1. Pause score difference were found within the control and HD replicates.

**fig S7.**
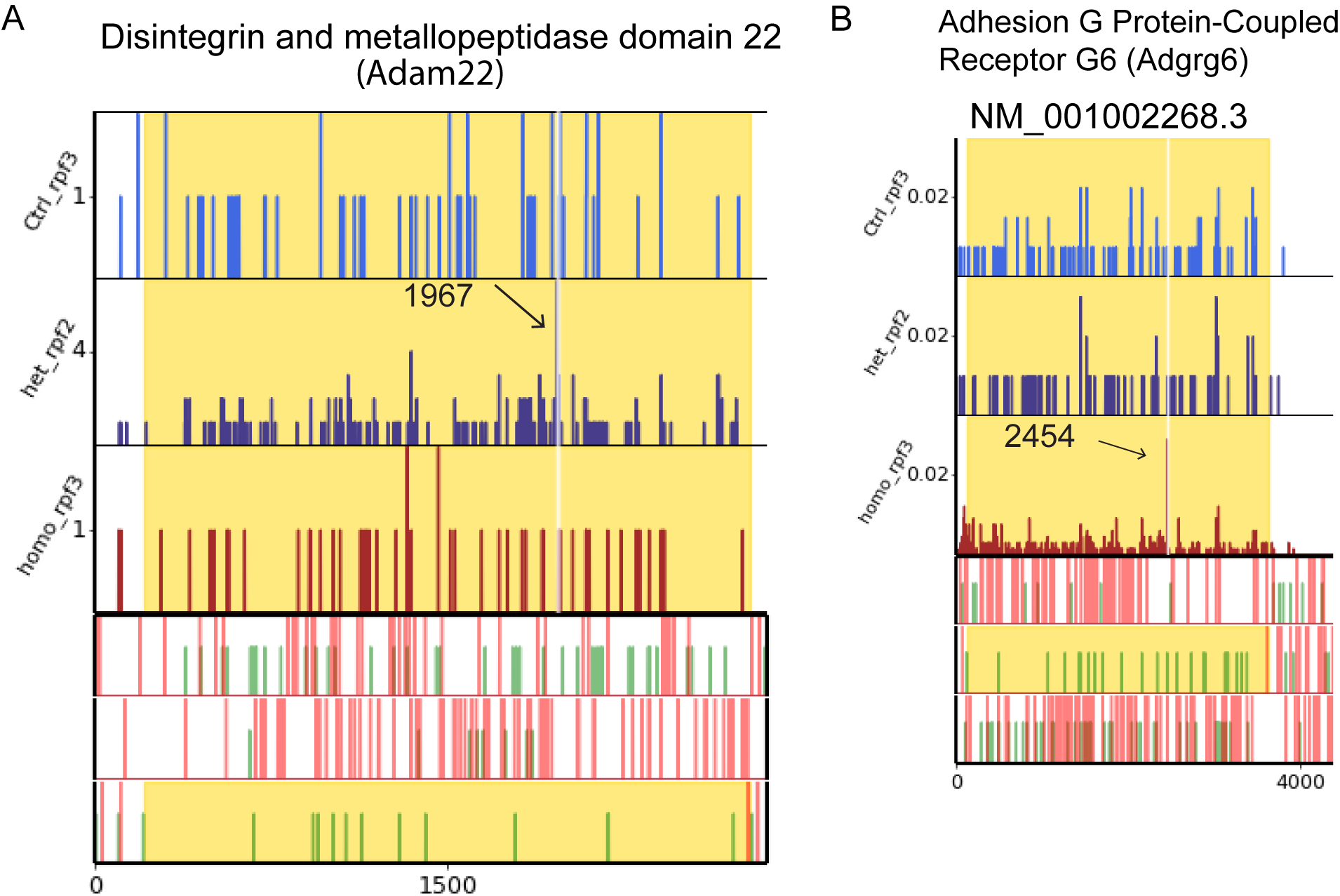
Single codon pause indicated with an arrow in *Adam22* in HD-Het (A) and *Adgrg6* in HD-homo (B).

**fig S8.**
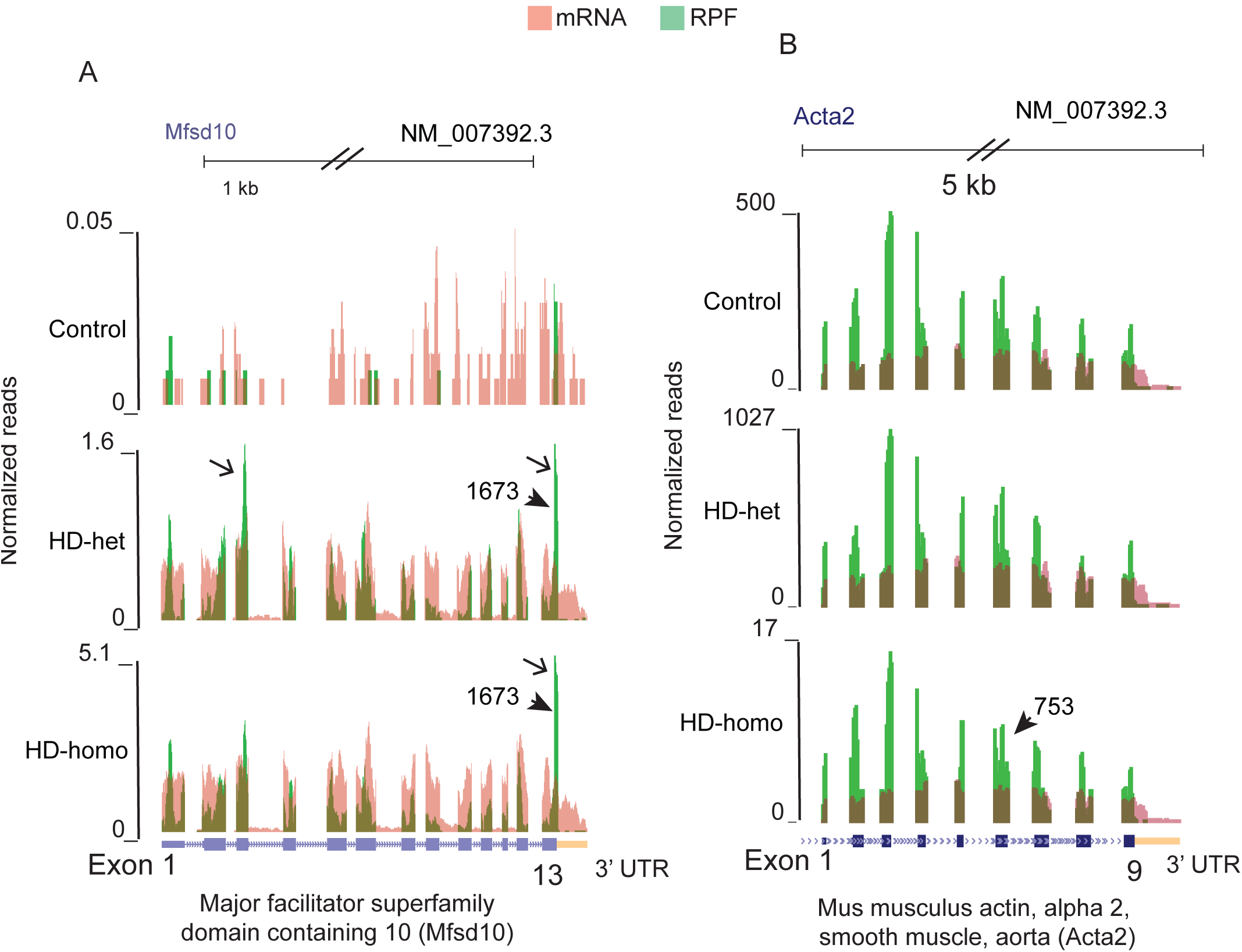
Normalized RPF/mRNA levels for A. *Mfsd10*, which showing a pause at 1673 (GAG, arrowhead) in exon 13 and higher RPF at 3’ and 5’ ends (arrow). B. *Acta2* showing pause location at 753 in exon 6 (CAG, arrowhead) and ∼15 fold less RPF compared to control.

**fig S9.**
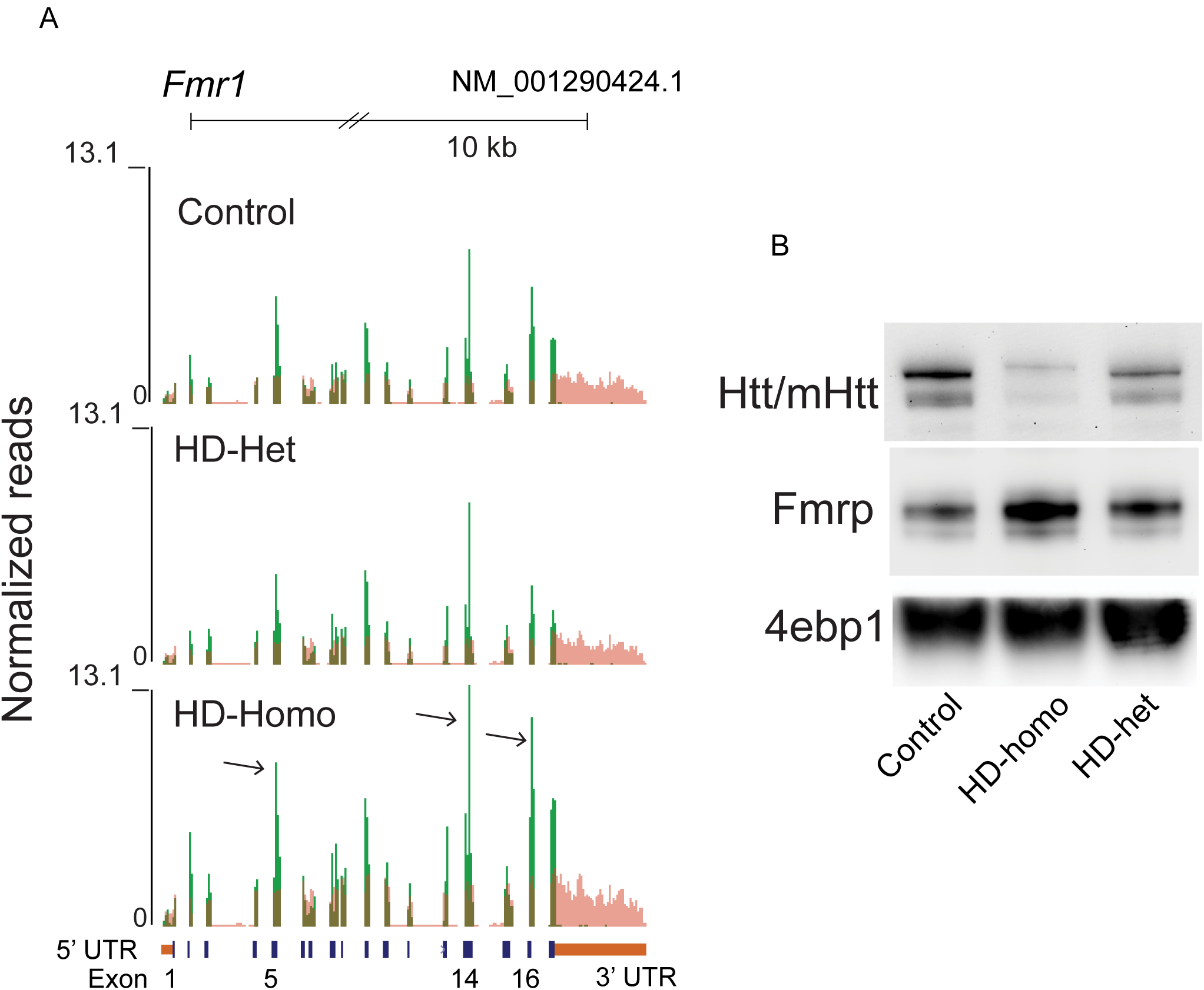
A. RPF and its accumulation on exons 5, 14, and 16 of *Fmr1* in HD-homo cells (arrows), compared to HD-het or control cells. B. Representative western blot showing indicated proteins in control, HD-homo ad HD-het cells.

**sfig S10.**
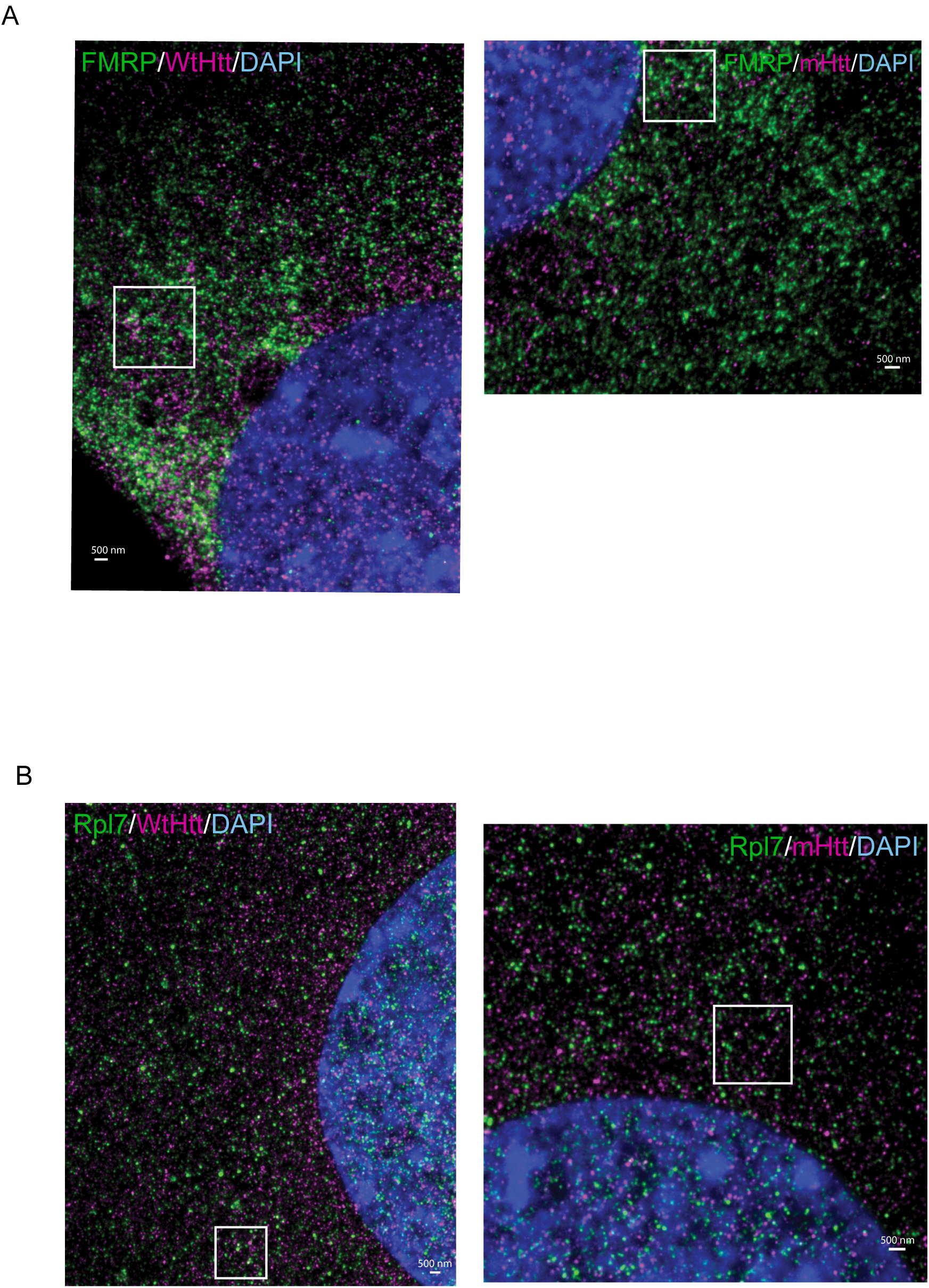
Representative STED image showing localization of FMRP/Htt or FMRP/mHtt in control and HD-homo cells (A) or Rpl7/Htt or Rpl7/mHtt in control and HD-homo cells (B), using immunochemical technique. Inset indicate the areas shown in main figures 4 and 6. DAPI, nuclear stain.

**fig S11.**
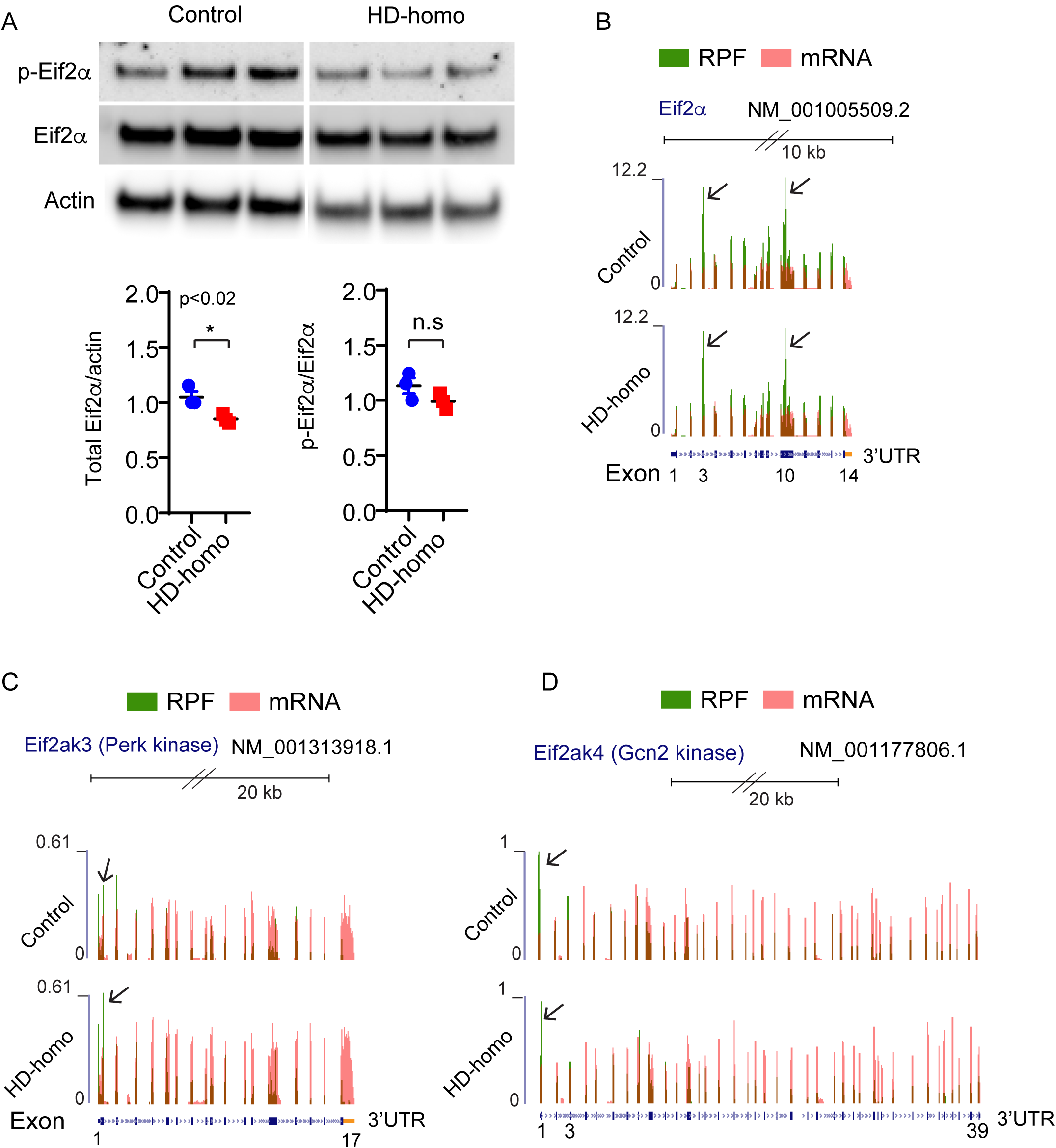
Representative Western blot showing phosphorylation of Eif2α (S51) and total Eif2α protein and their quantification in control and HD-homo cells. (B-D) Show RPF distribution in *Eif2a* (B), *Eif2ak3* (Perk kinase, C) or *Eif2ak4* (Gcn2 kinase, D) transcripts in control and HD-homo cells.

**fig S12.**
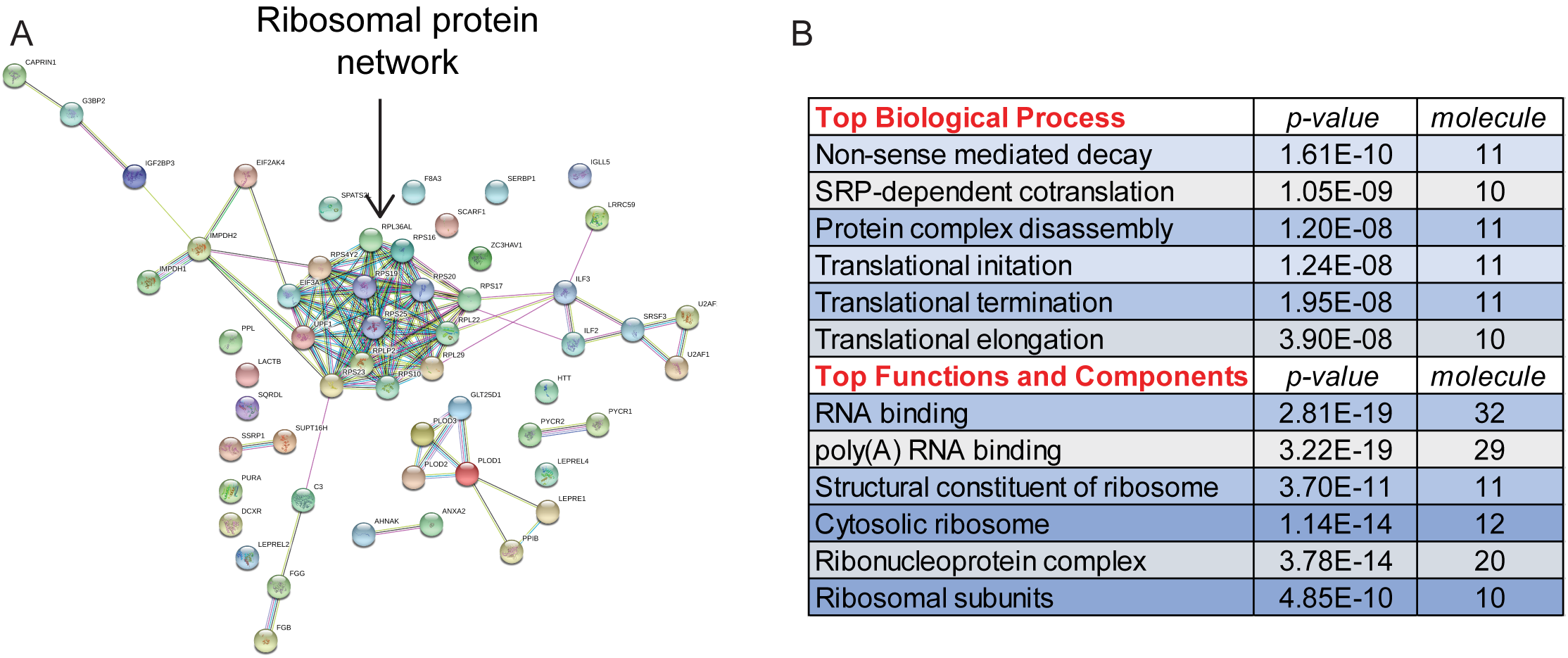
STRING/IPA analysis found translation and ribosome/RNA binding components (A), and an enrichment of the biological process related to ribosomes and RNA (B) from IP/MS/MS of HTT/mHTT IP from human fibroblasts, control (23Q/-) and HD (69Q/-).

**fig S13.**
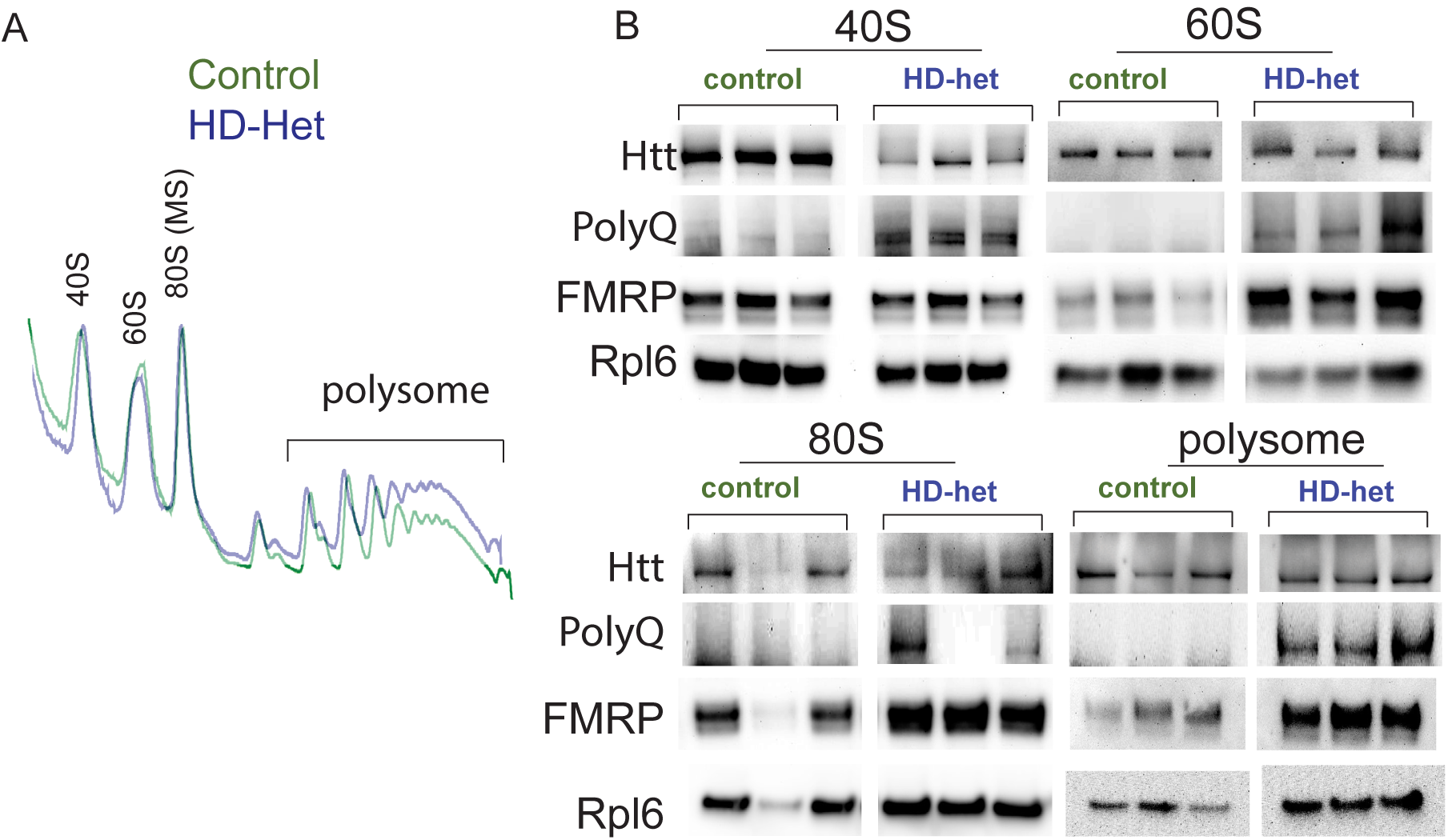
A. Represenetative ribosome fractionations of striatal cells. B. Western blot analysis of indicated proteins in different fractions.

**fig S14.**
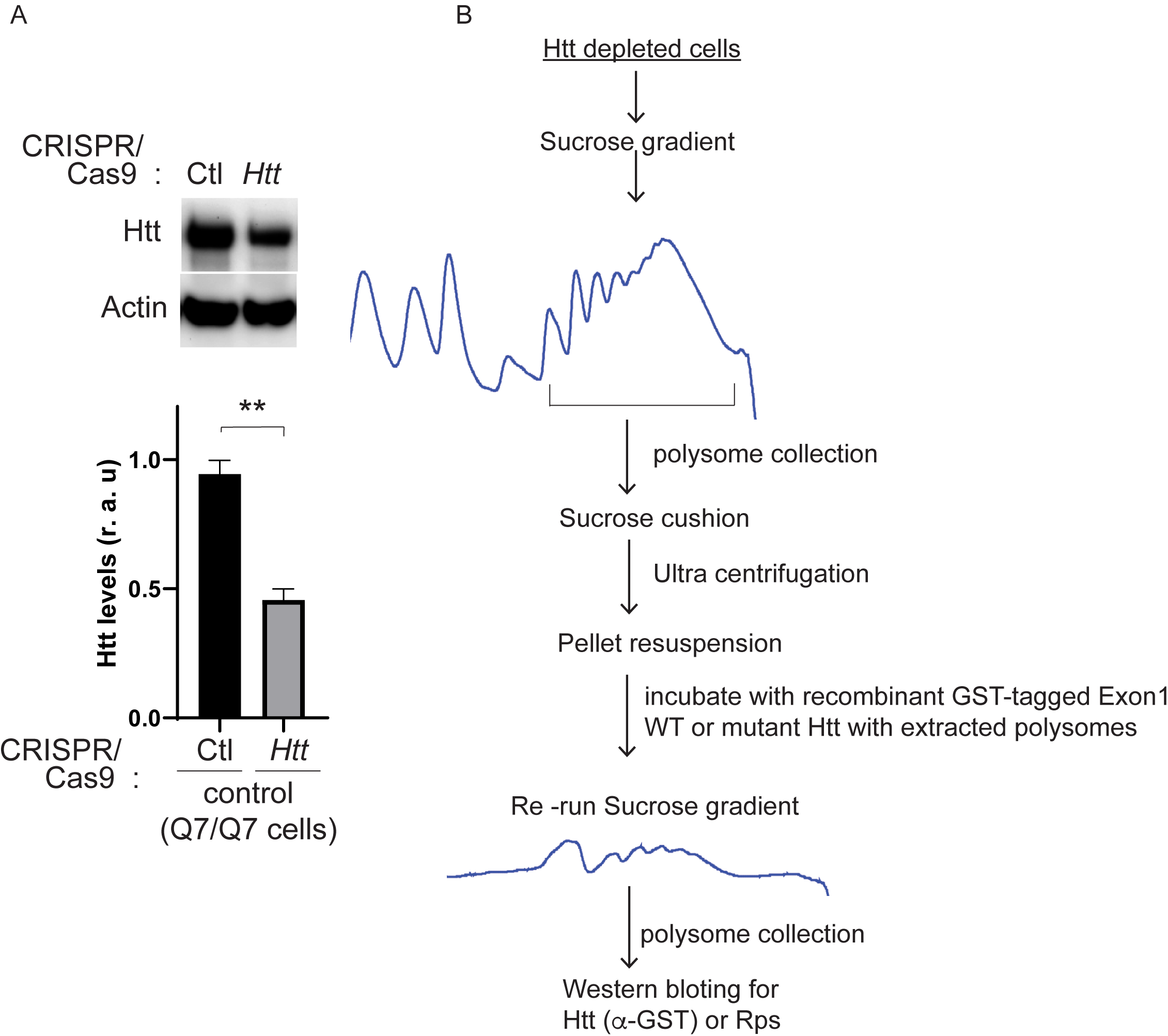
A. Represenetative Western blot showing indicated protein and Htt quantification (lower panel). B. Experimental plan for mHTT in vitro ribosome binding assay in Htt depleted cells.

**fig S15.**
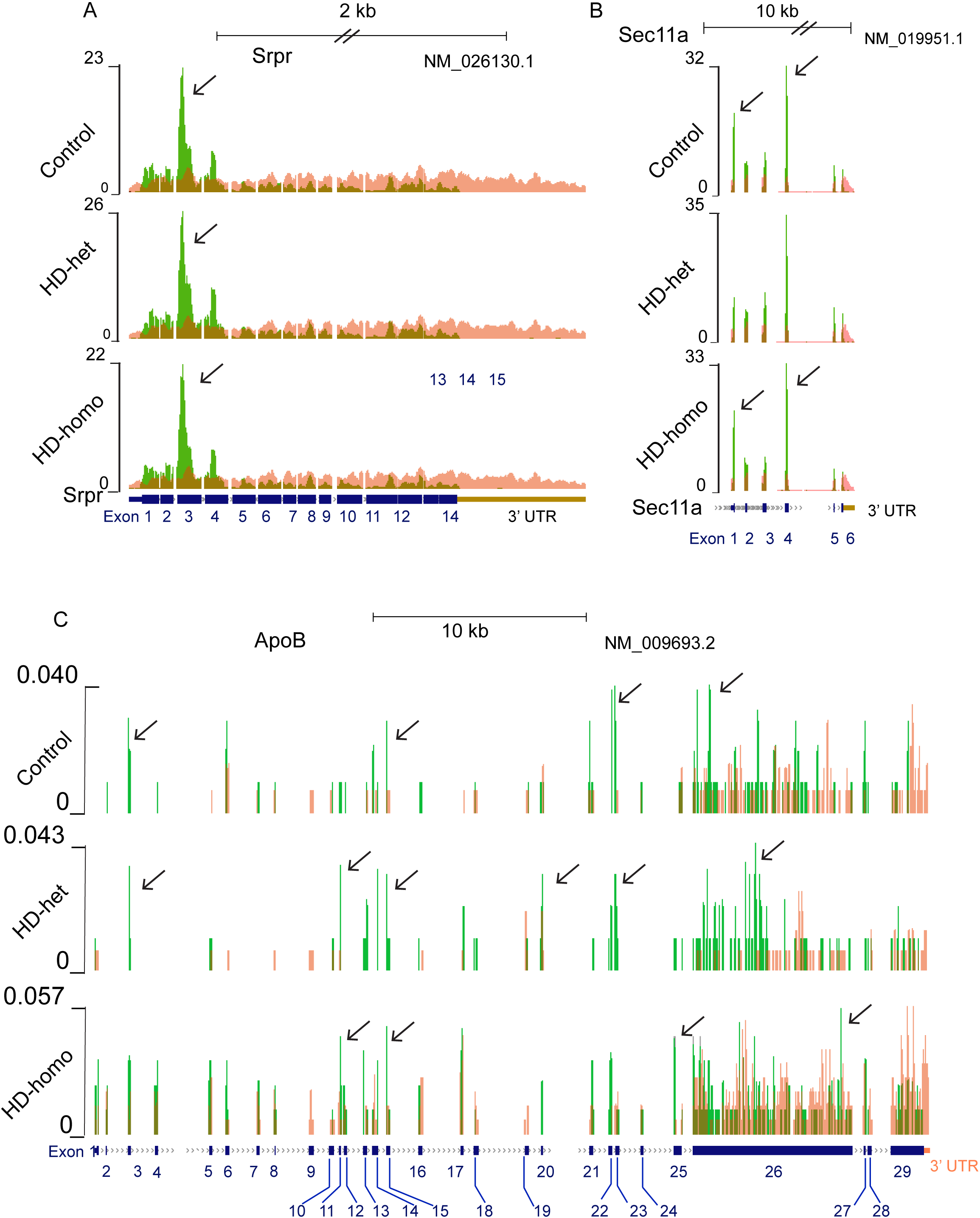
RPF/mRNA profile of indicated transcript (A) *Srpr* show ribosome occupany at exon 3 (arrow). (B) *Sec11* show occupancy at exon 1 and 4 (arrow). (C) *Apo B*, differential ribosome occupancy at various exons (arrow).

## REFERENCES

1. M. P. Duyao et al., Inactivation of the mouse Huntington’s disease gene homolog Hdh. Science 269, 407–410 (1995).

2. J. Nasir et al., Targeted disruption of the Huntington’s disease gene results in embryonic lethality and behavioral and morphological changes in heterozygotes. Cell 81, 811–823 (1995).

3. S. Zeitlin, J. P. Liu, D. L. Chapman, V. E. Papaioannou, A. Efstratiadis, Increased apoptosis and early embryonic lethality in mice nullizygous for the Huntington’s disease gene homologue. Nat Genet 11, 155–163 (1995).

4. J. K. White et al., Huntingtin is required for neurogenesis and is not impaired by the Huntington’s disease CAG expansion. Nat Genet 17, 404–410 (1997).

5. G. Wang, X. Liu, M. A. Gaertig, S. Li, X. J. Li, Ablation of huntingtin in adult neurons is nondeleterious but its depletion in young mice causes acute pancreatitis. Proc Natl Acad Sci U S A 113, 3359–3364 (2016).

6. I. Dragatsis et al., Effect of early embryonic deletion of huntingtin from pyramidal neurons on the development and long-term survival of neurons in cerebral cortex and striatum. Neurobiol Dis 111, 102–117 (2018).

7. J. P. Liu, S. O. Zeitlin, Is Huntingtin Dispensable in the Adult Brain? J Huntingtons Dis 6, 1–17 (2017).

8. R. H. Myers et al., Homozygote for Huntington disease. Am J Hum Genet 45, 615–618 (1989).

9. N. S. Wexler et al., Homozygotes for Huntington’s disease. Nature 326, 194–197 (1987).

10. M. DiFiglia et al., Huntingtin is a cytoplasmic protein associated with vesicles in human and rat brain neurons. Neuron 14, 1075–1081 (1995).

11. G. Hoffner, P. Kahlem, P. Djian, Perinuclear localization of huntingtin as a consequence of its binding to microtubules through an interaction with beta-tubulin: relevance to Huntington’s disease. J Cell Sci 115, 941–948 (2002).

12. J. Velier et al., Wild-type and mutant huntingtins function in vesicle trafficking in the secretory and endocytic pathways. Exp Neurol 152, 34–40 (1998).

13. H. Brandstaetter, A. J. Kruppa, F. Buss, Huntingtin is required for ER-to-Golgi transport and for secretory vesicle fusion at the plasma membrane. Dis Model Mech 7, 1335–1340 (2014).

14. J. Xia, D. H. Lee, J. Taylor, M. Vandelft, R. Truant, Huntingtin contains a highly conserved nuclear export signal. Hum Mol Genet 12, 1393–1403 (2003).

15. K. B. Kegel et al., Huntingtin is present in the nucleus, interacts with the transcriptional corepressor C-terminal binding protein, and represses transcription. J Biol Chem 277, 7466–7476 (2002).

16. C. Zuccato et al., Loss of huntingtin-mediated BDNF gene transcription in Huntington’s disease. Science 293, 493–498 (2001).

17. S. Sipione et al., Early transcriptional profiles in huntingtin-inducible striatal cells by microarray analyses. Hum Mol Genet 11, 1953–1965 (2002).

18. F. C. Nucifora, Jr. et al., Interference by huntingtin and atrophin-1 with cbp-mediated transcription leading to cellular toxicity. Science 291, 2423–2428 (2001).

19. H. Zhang et al., Elucidating a normal function of huntingtin by functional and microarray analysis of huntingtin-null mouse embryonic fibroblasts. BMC Neurosci 9, 38 (2008).

20. H. Takano, J. F. Gusella, The predominantly HEAT-like motif structure of huntingtin and its association and coincident nuclear entry with dorsal, an NF-kB/Rel/dorsal family transcription factor. BMC Neurosci 3, 15 (2002).

21. J. S. Steffan et al., The Huntington’s disease protein interacts with p53 and CREB-binding protein and represses transcription. Proc Natl Acad Sci U S A 97, 6763–6768 (2000).

22. N. Neelagandan et al., TDP-43 enhances translation of specific mRNAs linked to neurodegenerative disease. Nucleic Acids Res 47, 341–361 (2019).

23. M. Kapur, C. E. Monaghan, S. L. Ackerman, Regulation of mRNA Translation in Neurons-A Matter of Life and Death. Neuron 96, 616–637 (2017).

24. W. M. Pryor et al., Huntingtin promotes mTORC1 signaling in the pathogenesis of Huntington’s disease. Sci Signal 7, ra103 (2014).

25. F. Trettel et al., Dominant phenotypes produced by the HD mutation in STHdh(Q111) striatal cells. Hum Mol Genet 9, 2799–2809 (2000).

26. E. K. Schmidt, G. Clavarino, M. Ceppi, P. Pierre, SUnSET, a nonradioactive method to monitor protein synthesis. Nat Methods 6, 275–277 (2009).

27. N. T. Ingolia, L. F. Lareau, J. S. Weissman, Ribosome profiling of mouse embryonic stem cells reveals the complexity and dynamics of mammalian proteomes. Cell 147, 789–802 (2011).

28. X. Yan, T. A. Hoek, R. D. Vale, M. E. Tanenbaum, Dynamics of Translation of Single mRNA Molecules In Vivo. Cell 165, 976–989 (2016).

29. N. T. Ingolia, S. Ghaemmaghami, J. R. Newman, J. S. Weissman, Genome-wide analysis in vivo of translation with nucleotide resolution using ribosome profiling. Science 324, 218–223 (2009).

30. G. A. Brar et al., High-resolution view of the yeast meiotic program revealed by ribosome profiling. Science 335, 552–557 (2012).

31. S. Zhang et al., Analysis of Ribosome Stalling and Translation Elongation Dynamics by Deep Learning. Cell Syst 5, 212–220 e216 (2017).

32. J. K. Kim, M. J. Hollingsworth, Localization of in vivo ribosome pause sites. Anal Biochem 206, 183–188 (1992).

33. B. Liu et al., Regulatory discrimination of mRNAs by FMRP controls mouse adult neural stem cell differentiation. Proc Natl Acad Sci U S A 115, E11397–E11405 (2018).

34. J. L. Christian Oertlin, Valentina Gandin, Carl Murie, Laia Masvidal, Marie Cargnello, Luc Furic, Ivan Topisirovic, Ola Larsson, Generally applicable transcriptome-wide analysis of translational efficiency using anota2seq. BioRxiv (Preprint), (May 14, 2018).

35. O. Larsson, N. Sonenberg, R. Nadon, Identification of differential translation in genome wide studies. Proc Natl Acad Sci U S A 107, 21487–21492 (2010).

36. J. H. Cha, Transcriptional dysregulation in Huntington’s disease. Trends Neurosci 23, 387–392 (2000).

37. M. Haeussler et al., The UCSC Genome Browser database: 2019 update. Nucleic Acids Res 47, D853–D858 (2019).

38. R. Kumari, A. M. Michel, P. V. Baranov, PausePred and Rfeet: webtools for inferring ribosome pauses and visualizing footprint density from ribosome profiling data. RNA 24, 1297–1304 (2018).

39. D. E. Andreev et al., Translation of 5’ leaders is pervasive in genes resistant to eIF2 repression. Elife 4, e03971 (2015).

40. B. Hivert, L. Marien, K. N. Agbam, C. Faivre-Sarrailh, ADAM22 and ADAM23 modulate the targeting of the Kv1 channel-associated protein LGI1 to the axon initial segment. J Cell Sci 132, (2019).

41. J. C. Darnell, E. Klann, The translation of translational control by FMRP: therapeutic targets for FXS. Nat Neurosci 16, 1530–1536 (2013).

42. M. Ascano, Jr. et al., FMRP targets distinct mRNA sequence elements to regulate protein expression. Nature 492, 382–386 (2012).

43. M. Shen et al., Reduced mitochondrial fusion and Huntingtin levels contribute to impaired dendritic maturation and behavioral deficits in Fmr1-mutant mice. Nat Neurosci 22, 386–400 (2019).

44. T. D. Baird, R. C. Wek, Eukaryotic initiation factor 2 phosphorylation and translational control in metabolism. Adv Nutr 3, 307–321 (2012).

45. A. L. Southwell et al., An enhanced Q175 knock-in mouse model of Huntington disease with higher mutant huntingtin levels and accelerated disease phenotypes. Hum Mol Genet 25, 3654–3675 (2016).

46. B. P. Culver et al., Proteomic analysis of wild-type and mutant huntingtin-associated proteins in mouse brains identifies unique interactions and involvement in protein synthesis. J Biol Chem 287, 21599–21614 (2012).

47. T. Ratovitski et al., Huntingtin protein interactions altered by polyglutamine expansion as determined by quantitative proteomic analysis. Cell Cycle 11, 2006–2021 (2012).

48. E. Scherzinger et al., Self-assembly of polyglutamine-containing huntingtin fragments into amyloid-like fibrils: implications for Huntington’s disease pathology. Proc Natl Acad Sci U S A 96, 4604–4609 (1999).

49. C. V. Nicchitta, T. Zheng, Regulation of the ribosome-membrane junction at early stages of presecretory protein translocation in the mammalian endoplasmic reticulum. J Cell Biol 139, 1697–1708 (1997).

50. A. Protzel, A. J. Morris, Gel chromatographic analysis of nascent globin chains. Evidence of nonuniform size distribution. J Biol Chem 249, 4594–4600 (1974).

51. D. E. Andreev et al., Insights into the mechanisms of eukaryotic translation gained with ribosome profiling. Nucleic Acids Res 45, 513–526 (2017).

52. J. C. Young, D. W. Andrews, The signal recognition particle receptor alpha subunit assembles co-translationally on the endoplasmic reticulum membrane during an mRNA-encoded translation pause in vitro. EMBO J 15, 172–181 (1996).

53. M. H. Kivlen, C. A. Dorsey, V. R. Lingappa, R. S. Hegde, Asymmetric distribution of pause transfer sequences in apolipoprotein B-100. J Lipid Res 38, 1149–1162 (1997).

54. M. J. Hollingsworth, J. K. Kim, N. E. Stollar, Heelprinting analysis of in vivo ribosome pause sites. Methods Mol Biol 77, 153–165 (1998).

55. J. C. Darnell et al., FMRP stalls ribosomal translocation on mRNAs linked to synaptic function and autism. Cell 146, 247–261 (2011).

56. W. T. Greenough et al., Synaptic regulation of protein synthesis and the fragile X protein. Proc Natl Acad Sci U S A 98, 7101–7106 (2001).

57. L. K. Myrick et al., Independent role for presynaptic FMRP revealed by an FMR1 missense mutation associated with intellectual disability and seizures. Proc Natl Acad Sci U S A 112, 949–956 (2015).

58. F. Bedogni et al., Autism susceptibility candidate 2 (Auts2) encodes a nuclear protein expressed in developing brain regions implicated in autism neuropathology. Gene Expr Patterns 10, 9–15 (2010).

59. S. Dowler et al., Identification of pleckstrin-homology-domain-containing proteins with novel phosphoinositide-binding specificities. Biochem J 351, 19–31 (2000).

60. G. Allavena et al., Suppressed translation as a mechanism of initiation of CASP8 (caspase 8)-dependent apoptosis in autophagy-deficient NSCLC cells under nutrient limitation. Autophagy 14, 252–268 (2018).

61. L. M. Knowles, C. Yang, A. Osterman, J. W. Smith, Inhibition of fatty-acid synthase induces caspase-8-mediated tumor cell apoptosis by up-regulating DDIT4. J Biol Chem 283, 31378–31384 (2008).

62. M. Flasza, P. Gorman, R. Roylance, A. E. Canfield, M. Baron, Alternative splicing determines the domain structure of WWP1, a Nedd4 family protein. Biochem Biophys Res Commun 290, 431–437 (2002).

63. S. Subramaniam, K. M. Sixt, R. Barrow, S. H. Snyder, Rhes, a striatal specific protein, mediates mutant-huntingtin cytotoxicity. Science 324, 1327–1330 (2009).

64. M. M, Cutadapt Removes Adapter Sequences From High-Throughput Sequencing Reads. EMBnet.journal 17 (I), 10–12 (2011).

65. B. Li, C. N. Dewey, RSEM: accurate transcript quantification from RNA-Seq data with or without a reference genome. BMC Bioinformatics 12, 323 (2011).

66. A. R. Quinlan, I. M. Hall, BEDTools: a flexible suite of utilities for comparing genomic features. Bioinformatics 26, 841–842 (2010).

67. M. I. Love, W. Huber, S. Anders, Moderated estimation of fold change and dispersion for RNA-seq data with DESeq2. Genome Biol 15, 550 (2014).

68. N. Shahani et al., RasGRP1 promotes amphetamine-induced motor behavior through a Rhes interaction network (“Rhesactome”) in the striatum. Sci Signal 9, ra111 (2016).

